# Assigning Targetable Pathways to Transdiagnostic Subgroups Across Neurodevelopmental Disorders

**DOI:** 10.1101/2025.03.04.641443

**Authors:** Jacob Ellegood, Antoine Beauchamp, Yohan C. Yee, Gabriel A. Devenyi, Justine Ziolkowski, Lily R. Qiu, Rand Askalan, Muhammad Ayub, Phillip Suetterlin, Alex P.A. Donovan, Leticia Pérez-Sisqués, M. Albert Basson, Katherine Quesnel, Nathalie G. Berube, Taeseon Woo, David Beversdorf, Hans Bjornsson, Randy Blakely, Jacqueline Crawley, Jennifer Crosbie, Brian O. Orr, Graeme Davis, Matthew Genestine, Emanuel DiCicco-Bloom, Sean Egan, Kyle D. Fink, Sarah Asbury, Jonathan Lai, Kelly Rilett, Jane Foster, John B. Vincent, Paul W. Frankland, Stelios Georgiades, Olga Penagarikano, Daniel Geschwind, Roman J. Giger, Sander Markx, Joseph Gogos, Christelle Golzio, Rafaela Goveia, Marco Pagani, Alessandro Gozzi, Laura K Pacey, David R. Hampson, Tzyy-Nan Huang, Tzu-Li Yen, Yi-Ping Hsueh, Alana Iaboni, Megha Amar, Lilia M Iakoucheva, Jessica K Jones, Elizabeth A Kelley, Brigette Kieffer, Mihyun Bae, Hwajin Jung, Hyosang Kim, Haram Park, Junye Daniel Roh, Eunjoon Kim, Genevieve Konopka, Christine Laliberté, Julie Lefebvre, Kathie Eagleson, Pat Levitt, Aurea B Martins-Bach, Thomas J. Cunningham, Elizabeth Fisher, Karla L. Miller, Alea A. Mills, Alysson R. Muotri, Rob Nicolson, Leigh Spencer Noakes, Brian J. Nieman, Cesar Canales, Alexander S. Nord, Lauryl M. J. Nutter, E Tam, Lucy R. Osborne, Amy Clipperton-Allen, Damon Page, Brooke Babineau, Hyang Mi Moon, Theo Palmer, Keqin Yan, David J. Picketts, Qiangqiang Xia, Craig M. Powell, Armin Raznahan, Diane M. Robins, Gavin Rumbaugh, Ameet S. Sengar, Michael W. Salter, Russell Schachar, Lia D’Abate, Clarrisa A. Bradley, Sang Yoon Ko, Stephen W. Scherer, Nycole A. Copping, Stela P. Pektova, Jill Silverman, Karun Singh, Nam-Shik Kim, Ki-Jun Yoon, Guo-li Ming, Hongjun Song, Shoshana Spring, Jin Nakatani, N Nakai, Jun Nomura, Toru Takumi, Peter Tsai, Matthew Bruce, Karen Jones, Judy Van de Water, Matthijs C. van Eede, Travis M Kerr, Chris Muller, Jeremy Veenstra-VanderWeele, Marlee Vandewouw, Rosanna Weksberg, Rachel Wevrick, Haim Belinson, Anthony Wynshaw-Boris, Konstantinos Zarbalis, Brett Trost, Margot J. Taylor, Rogier B. Mars, M. Mallar Chakravarty, Azadeh Kushki, Evdokia Anagnostou, Jason P. Lerch

**Affiliations:** Mouse Imaging Centre, Hospital for Sick Children, Toronto, Ontario, Canada; Holland Bloorview Kids Rehabilitation Hospital, Toronto, Ontario, Canada; Cumming School of Medicine, Department of Radiology, University of Calgary, Calgary, Alberta, Canada; The Douglas Research Centre, McGill University, Montreal, Quebec, Canada; Oxford Centre for Integrative Neuroimaging, University of Oxford, Oxford, UK; Queen’s University, Kingston, Ontario, Canada; Centre for Craniofacial and Regenerative Biology, King’s College London, London, UK; LMB, Cambridge, UK; MRC Centre for Neurodevelopmental Disorders, King’s College London, London, UK; Clinical and Biomedical Sciences, University of Exeter Medical School, Exeter, UK; Departments of Anatomy & Cell Biology and Paediatrics, Western University, London, Canada; Division of Genetics & Development, Children’s Health Research Institute, London ON, Canada; Departments of Anatomy & Cell Biology, Paediatrics, and Oncology, Western University, London, Canada; Department of Molecular Pathobiology, College of Dentistry, New York University, New York, NY; Departments of Neurology, Radiology, and Psychological Sciences, University of Missouri, William and Nancy Thompson Endowed Chair in Radiology; The Louma G. Laboratory of Epigenetic Research, Faculty of Medicine, University of Iceland, Reykjavik, Iceland; Department of Genetics and Molecular Medicine, Landspitali University Hospital, Reykjavik, Iceland; McKusick-Nathans Department of Genetic Medicine, Johns Hopkins University School of Medicine; Baltimore, MD, USA; Department of Biomedical Science, Charles E. Schmidt College of Medicine, Jupiter, FL, USA; Stiles-Nicholson Brain Institute, Florida Atlantic University, Jupiter, FL, USA; MIND Institute, University of California Davis School of Medicine, Sacramento, CA, USA; Program in Neuroscience & Mental Health, The Hospital for Sick Children, Toronto, Ontario, Canada; Department of Biochemistry and Biophysics, University of California, San Francisco, CA, USA; Neuroscience and Cell Biology/Pediatrics, Rutgers Robert Wood Johnson Medical School, NJ, USA; Program in Cell & Systems Biology, The Hospital for Sick Children, Toronto, Ontario, Canada; Department of Psychiatry & Behavioural Neurosciences, McMaster University, Hamilton, ON, Canada; Institute of Health Policy, Management and Evaluation, University of Toronto, Toronto, Ontario, Canada; Department of Psychiatry, UT Southwestern, Dallas, TX, USA; Center for Depression Research and Clinical Care, UT Southwestern, Dallas, TX, USA; O’Donnell Brain Institute, UT Southwestern, Dallas, TX, USA; Molecular Brain Science, Centre for Addiction & Mental Health, Toronto, Ontario, Canada; Department of Psychiatry, University of Toronto, Toronto, Ontario, Canada; Institute of Medical Science, University of Toronto, Toronto, Ontario, Canada; Department of Psychology, University of Toronto, Toronto, Ontario, Canada; Department of Physiology, University of Toronto, Toronto, Ontario, Canada; David Geffen School of Medicine, UCLA, LA, CA, USA; Department of Cell and Developmental Biology, University of Michigan Medical School, Ann Arbor MI, USA; Department of Neurology, University of Michigan Medical School, Ann Arbor, MI, USA; Mortimer B. Zuckerman Mind Brain and Behavior Institute, Columbia University, New York, NY, USA; Stavros Niarchos Foundation Center for Precision Psychiatry and Mental Health, Columbia University Irving Medical Center, New York, NY, USA; Université de Strasbourg, CNRS, Inserm, IGBMC UMR 7104-UMR-S 1258, F-67404 Illkirch, France; Functional Neuroimaging Laboratory, Istituto Italiano di Tecnologia, Rovereto, Italy; Department of Pharmaceutical Sciences, University of Toronto, Toronto, Ontario, Canada; Institute of Molecular Biology, Academia Sinica, Taipei, 11529, Taiwan, ROC; Department of Psychiatry, University of California San Diego, La Jolla, CA, USA; Center for Synaptic Brain Dysfunctions, Institute for Basic Science (IBS), Daejeon 34141, Korea; Department of Neuroscience, Peter O’Donnell Jr. Brain Institute, UT Southwestern Medical Center, Dallas, TX, USA; Children’s Hospital Los Angeles, LA, CA, USA; Cold Spring Harbor Laboratory, Laurel Hollow, NY, USA; Department of Pediatrics and Cellular & Molecular Medicine, School of Medicine, University of California San Diego, La Jolla, CA 92093, USA; Department of Psychiatry, University of Western Ontario, London, Ontario, Canada; Program in Translational Medicine, The Hospital for Sick Children, Toronto, ON, Canada; Program in Genetics and Genome Biology, The Hospital for Sick Children, Toronto, ON, Canada; The Centre for Phenogenomics, Toronto, ON, Canada; Departments of Medicine and Molecular Genetics, University of Toronto, Toronto, ON, Canada; Department of Pediatrics, University of Washington & Norcliffe Foundation Center for Integrative Brain Research, Seattle Children’s Research Institute, Seattle, WA; Department of Neurosurgery, Institute for Stem Cell Biology and Regenerative Medicine, Stanford University, School of Medicine, Stanford, CA, USA; Regenerative Medicine Program, Ottawa Hospital Research Institute, Ottawa, ON, CANADA; Departments of Medicine, Cellular & Molecular Medicine, and Biochemistry, Microbiology & Immunology, University of Ottawa, Ottawa, ON, CANADA; Department of Neurobiology & Civitan International Research Center, University of Alabama at Birmingham Heersink School of Medicine, Birmingham, AL, USA; Section on Developmental Neurogenomics, NIMH Intramural Research Program, Bethesda, MD, USA; Department of Human Genetics, University of Michigan, MI, USA; The Herbert Wertheim UF Scripps Institute for Biomedical Innovation & Technology, Jupiter, FL, USA; Program in Genetics and Genome Biology and The Centre for Applied Genomics, The Hospital for Sick Children; McLaughlin Centre and Department of Molecular Genetics, University of Toronto; Department of Neuroscience, University of Pennsylvania, Philadelphia, PA, USA; Department of Biomedical Sciences, College of Life Sciences, Ritsumeikan University, Shiga, Japan; Department of Physiology and Cell Biology, Kobe University School of Medicine, Chuo, Kobe, Japan; Departments of Neurology, Neuroscience, Psychiatry, and Pediatrics, University of Texas Southwestern Medical Center, Dallas, TX; Department of Psychiatry, Columbia University, New York, NY, USA; New York State Psychiatric Institute, New York, NY, USA; Diagnostic & Interventional Radiology, Hospital for Sick Children, Toronto, Ontario, Canada; Paediatrics and Medical Genetics, Hospital for Sick Children, Toronto, Ontario, Canada; Department Of Medical Genetics, University of Alberta, Edmonton, Alberta, Canada; Department of Genetics and Genome Sciences, Case Western Reserve University School of Medicine, Cleveland, OH, USA; Department of Pathology and Laboratory Medicine, University of California, Davis; Institute for Pediatric Regenerative Medicine, Shriners Hospitals for Children, Northern California, 2425 Stockton Boulevard, Sacramento, CA 95817; Donders Institute for Brain, Cognition and Behaviour, Radboud University Nijmegen, Nijmegen, The Netherlands

## Abstract

The heterogeneity of autism and related neurodevelopmental conditions has impeded accurate prognoses and treatment discovery. Using translational neuroimaging across 135 mouse models (3,515 mice) and two human MRI datasets (n = 1,234 and n = 1,015), we derived participant subgroups from shared neuroanatomical features. These subgroups did not distinguish autism, ADHD, or OCD diagnoses and only modestly differentiated cognitive and behavioural phenotypes. Instead, they mapped onto four molecular pathways: (1) synaptic function; (2) MAPK and Wnt signalling; (3) chromatin modification and cellular stress responses; and (4) broader chromatin, immune, and second-messenger signalling pathways. This framework bridges preclinical models and idiopathic human neurodevelopmental conditions, linking patients to biologically relevant molecular mechanisms.

## Background

Neurodevelopmental disorders (NDDs), including autism spectrum disorder (ASD), attention deficit hyperactivity disorder (ADHD), and obsessive compulsive disorder (OCD), are traditionally defined based on behavioral and cognitive characteristics that emerge during childhood. Diagnosis or diagnostic label is often provided based on care-giver, teacher, psychologist, or psychiatric evaluation independent of underlying biology. Thus, these definitions do not consider biological mechanisms and lead to a genetically, behaviourally, and neuroanatomically heterogeneous grouping even within the individual disorders. Advances in neuroimaging(*1–3*), genetics(*4–7*), and large scale transdiagnostic datasets(*8*, *9*) suggest that NDDs may share overlapping neural substrates and molecular pathways, which challenge the validity of the behaviourally- based classifications and may indicate a need to redefine how we classify NDDs. Moving forward, it remains critically important for prognosis and treatment to identify biologically informed transdiagnostic sub-groups within the NDD population that are more homogeneous.

The high heritability of NDDs indicates a robust genetic contribution to the etiology of these conditions(*10*, *11*), suggesting that genetics can be used to identify biologically relevant sub-populations. However, the landscape of genetic causation is far from straightforward, with rare *de novo* variants also contributing to etiology(*12*, *13*), and there are confounding effects of diverse environments. Even within autism, currently only ∼15-20% of autistic individuals have an identified causal variant(*14*), and no single gene accounts for more than 1% of the clinical population(*15–18*). The literature suggests a proportion of autism etiology can be attributed to multiple genes in addition to environmental risk factors(*15*). Since genetics alone cannot yet be used to fully sub-categorize the autistic population, there has been a search in recent years for phenotypically driven clusters. Subgrouping analyses based upon clinical features in autism have been conducted by multiple groups and primarily generate clusters based upon severity across domains. More recently, subgrouping based on techniques that include structural and/or functional neuroimaging data suggest that traditional diagnostic boundaries do not properly capture the neural diversity of NDDs(*2*, *19–22*).

Recent work has attempted to implicate mechanisms underlying each cluster by integrating publicly accessible gene expression datasets(*23*). The extent to which normative gene expression can explain pathophysiological mechanisms remains to be clarified. The biological underpinnings of these reported phenotypic clusters could be probed to an extent by augmenting the clusters with genetic information from the individuals; however, a very small number of participants share any one identifiable high-impact rare variant. Thus, mouse models may be used to investigate the association between NDD-related genes and a phenotype of interest, such as neuroanatomy. In a previous study, focused primarily on autism, we imaged 26 distinct autism-related genetic mouse models to identify clusters using whole-brain mesoscopic neuroanatomy(*24*). Similarly, Gozzi and colleagues have used functional imaging to assess and create autism related subtypes in mouse and human(*25*). The overarching goal of this project is to evaluate the similarity between the mouse models and human individuals, allowing us to directly translate molecular signatures from the mouse onto individual humans. To that end, we have expanded to include 135 different mouse models (3,515 individual mice) relevant to autism and related NDDs (table S1) and two human databases, with 1,234 and 1,015 human individuals with NDDs. Figure 1 displays an overview of the analysis pipeline (see fig S1 for more detail). In essence, by comparing the patterns of neuroanatomical morphology across this expansive set of mouse models, we were able to identify distinct neuroanatomical clusters in the mouse that illuminated both novel connections among the implicated genes and specific enrichment for distinct molecular pathways and therefore potential drug targets. Further, we were able to identify analogous neuroanatomically driven clusters in a human cohort of individuals with NDDs and typically-developing controls from the Province of Ontario Neurodevelopmental Disorders (POND) Network and an additional cohort provided to us by Dr. Margot Taylor(*26*) (POND+), and we replicated those findings in a separate dataset collected by the Healthy Brain Network (HBN), totalling 2,249 human participants. Lastly, using a common space approach based on the spatial expression patterns of mouse-human orthologous genes(*27*), we evaluate the similarity between the mouse and human clusters, allowing us to directly translate neuroanatomical clusters across species and map the molecular signatures of the mouse onto human subgroups. This approach allows us to bridge the gap between preclinical models and human disorders and link previously idiopathic human patients to pertinent genetics, molecular mechanisms, and pathways.

**Fig. 1.**
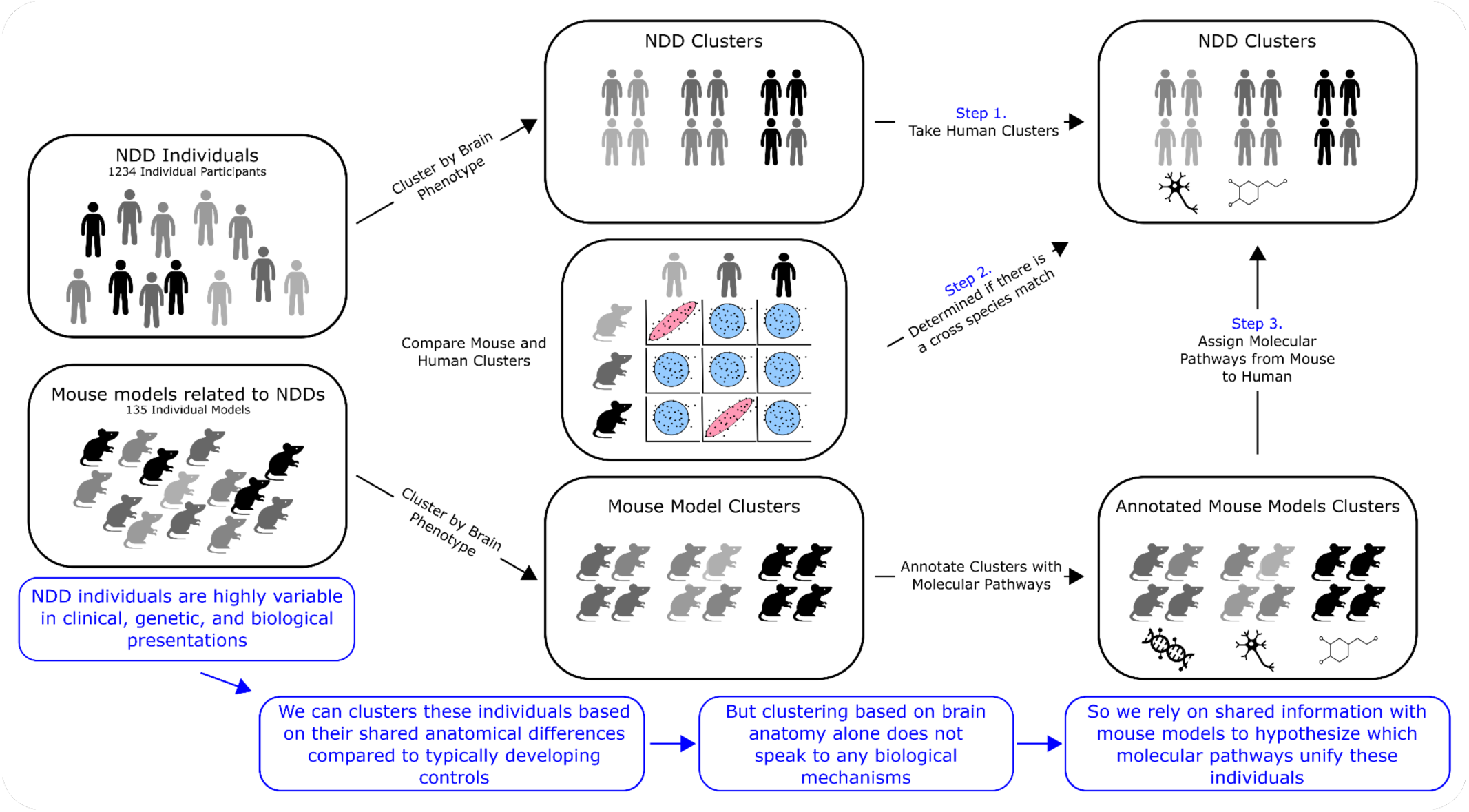
Overview of project workflow. On both the human and mouse sides of the analysis, we implemented an image processing workflow in which we used image registration to align the individual brain images to a species-specific common consensus average space, generated effect-size images for the human participants and mouse models, and separately identified clusters of human participants and mouse models based on neuroanatomical similarity. We evaluated the degree of similarity between the mouse and human clusters using freely available spatial gene expression data sets from the Allen Human Brain Atlas and Allen Mouse Brain Atlas. Mouse clusters were linked to molecular pathways using a combination of the STRING database (https://string-db.org)(28, *29*), publicly available gene lists the Bader group (https://www.baderlab.org)(30) and the Reactome database (https://reactome.org)(31). Following this workflow, we were able to link certain mouse and human clusters based on the similarity of their neuroanatomical morphology, and thus assign specific genetics and molecular pathway information directly to human participants.

## Results

We first wanted to understand how brain anatomy divided our mouse models into groups sharing common neuroanatomical phenotypes. Beginning with the effect size images, we constructed a weighted graph of the 135 models, which was then partitioned into clusters. An analysis of the clustering structure across possible solutions revealed robust evidence of organization at multiple scales in the similarity network (fig. S2), suggesting an interpretation of the cluster number (K) as a scale parameter for network organization. To narrow the scope of subsequent analyses, we focused on partitions with K = 2-10. At the largest scale in the network, the mouse models divided into clusters aligned along a cortical-cerebellar neural axis, with one end of this axis characterized by cortical increases and cerebellar decreases, and the other end featuring the reverse (fig. S3). As the clustering resolution is increased, the network is partitioned into groups of mouse models with distinct patterns of neuroanatomy, with clusters differentiating on anatomical patterns in midbrain, hindbrain, and cerebral nuclei in addition to the cortex-cerebellum axis (fig. S3, table S2). Overall, we found that brain anatomy separated the mouse models into distinct clusters in a multiscale fashion with a predominant axis of cortical-cerebellar variation.

We next examined how human participants with NDDs clustered based on their neuroanatomy, and whether these clusters aligned with those found in the mouse. Using the same pipeline, we performed network construction and clustering independently in the POND+ and HBN samples across the range K = 2-10 (fig. S4 and fig. S5). As with the mouse, we found that the participants in both samples broadly divide along a cortical-cerebellar axis at the largest scale. To quantify the association between mouse and human clusters, we projected them into a transcriptomic common space where they could be compared directly. Briefly, we constructed homologous gene expression signatures using spatial gene expression datasets from the Allen Institute for Brain Science, and evaluated the correlation between these signatures at multiple K. We find that the two clusters at the largest scale match across species, both in the POND+ and HBN datasets (fig. 2). Moreover, as the clustering resolution is increased, replicable cross-species matches can be found in 21 of the 54 mouse clusters. Of note are clusters of mouse models and human participants that persist across the solution set and consistently match across species (indicated by coloured strata in the Sankey diagram), highlighting robust and stable cross-species subgroups within the similarity networks. Overall, we found that clusters could directly be aligned in 39% of cases.

**Fig 2.**
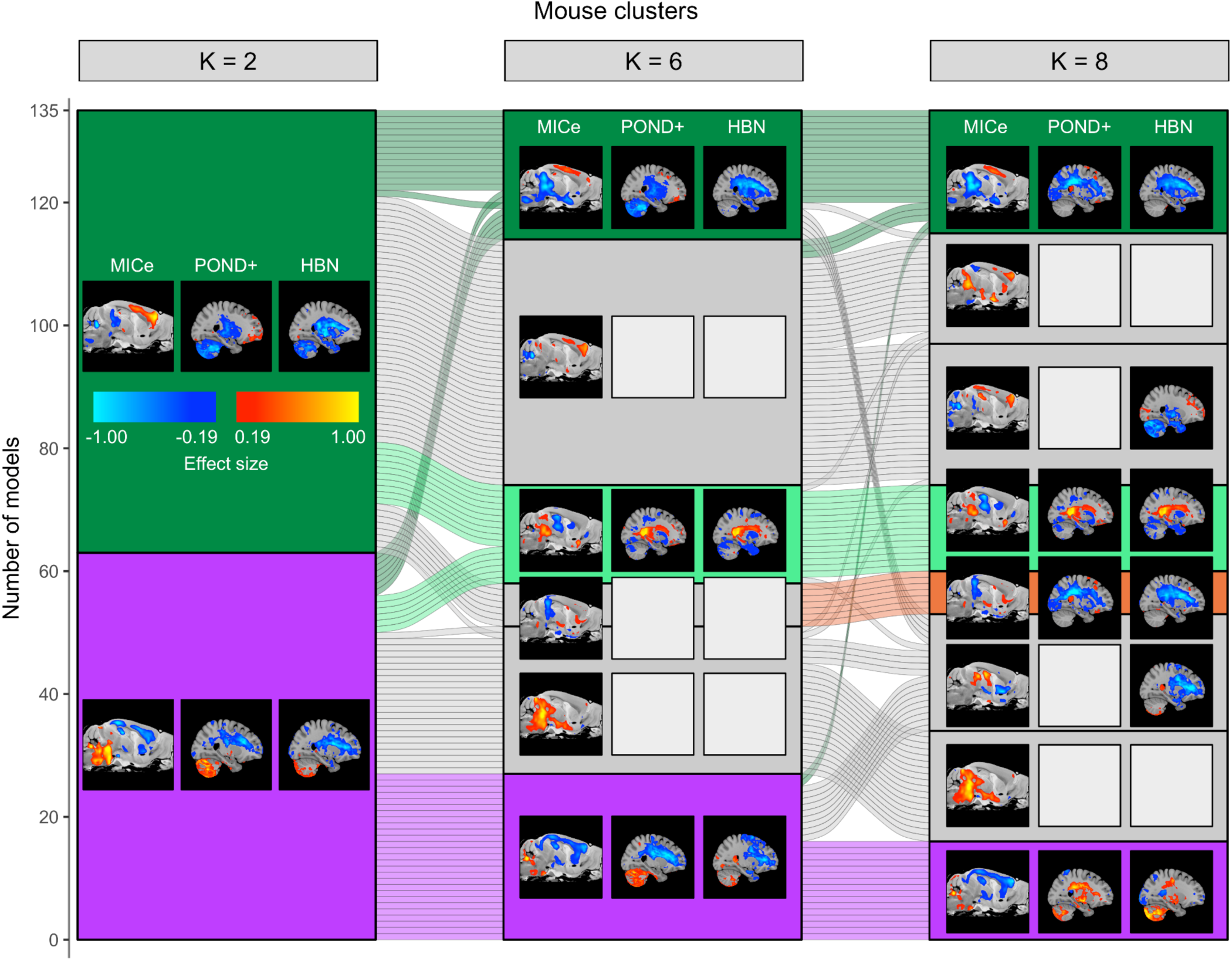
Mouse clusters define robust neuroanatomical groups with replicable human matches. Sankey diagram of mouse model clusters for K = 2, 6, 8. Matching human clusters in the POND+ and HBN samples are indicated where matches exist. Sagittal slices of the mouse and human cluster centroids are displayed. Empty squares are displayed in lieu of human slices when no matches exist. Mouse clusters with matches in both POND+ and HBN are coloured. These coloured groups indicate stable neuroanatomical subgroups across the clustering resolutions.

Having identified stable mouse clusters with robust correspondences in both POND+ and HBN, we then investigated whether these neuroanatomical subgroups were associated with unique molecular signalling pathways. Using the single gene models from our mouse clusters and their one-hop protein-protein interaction neighbours, we performed an enrichment analysis for molecular pathway modules in the Reactome database (table S3). At the largest scales in our network of mouse models, we find that the clusters are broadly enriched for most molecular pathways previously associated with neurodevelopmental disorders (fig. 3). However, as the clustering resolution is increased and the mouse models divide into subgroups with distinct neuroanatomy, we observe the emergence of increasingly specific molecular pathway signatures for those clusters. Focusing on four stable subgroups with matching clusters in the POND+ and HBN samples, we find a *MAPK-WNT group* (dark green) with enrichment for MAPK (average −log10(q) = 11.07), WNT (average −log10(q) = 22.21), receptor tyrosine kinases (average −log10(q) = 15.80), and GTPases (average −log10(q) = 11.78), which emerges at K = 6 and featuring prominent volume increases in isocortex and cerebellar cortex and volume decreases in the cerebral nuclei, midbrain, and hindbrain (fig. S6). We find a group composed only of the Androgen Receptor mouse models (*Ar group*, orange), enriched for signaling by nuclear receptors (average −log10(q) = 7.50), cellular responses to stress (average −log10(q) = 5.87), regulation of gene expression (average −log10(q) = 3.62), and chromatin modification (average −log10(q) = 3.78). We find a *synaptic-immune group* (light green), enriched for cytokine signaling in the immune system (average −log10(q) = 4.01), protein-protein interactions at synapses (average −log10(q) = 23.35), and transmission across chemical synapses (average −log10(q) = 13.65), featuring volume increase in the hind- and midbrain and volume decreases in the interbrain, cerebral nuclei, and hippocampus. Finally, we find a group that features low and diffuse enrichment across most of the solutions but narrows to transcription regulation (average −log10(q) = 4.48) and chromatin organization (average −log10(q) = 3.01) at the highest resolution (*transcription group*, purple), featuring large volume increases in the cerebellar cortex and hindbrain and volume decreases in isocortex and hippocampus. Together, these results demonstrate that clustering mouse models based on brain anatomy leads to translational subgroups that feature distinct molecular programs, with motifs such as MAPK-WNT, synaptic-immune, and transcription.

**Fig. 3.**
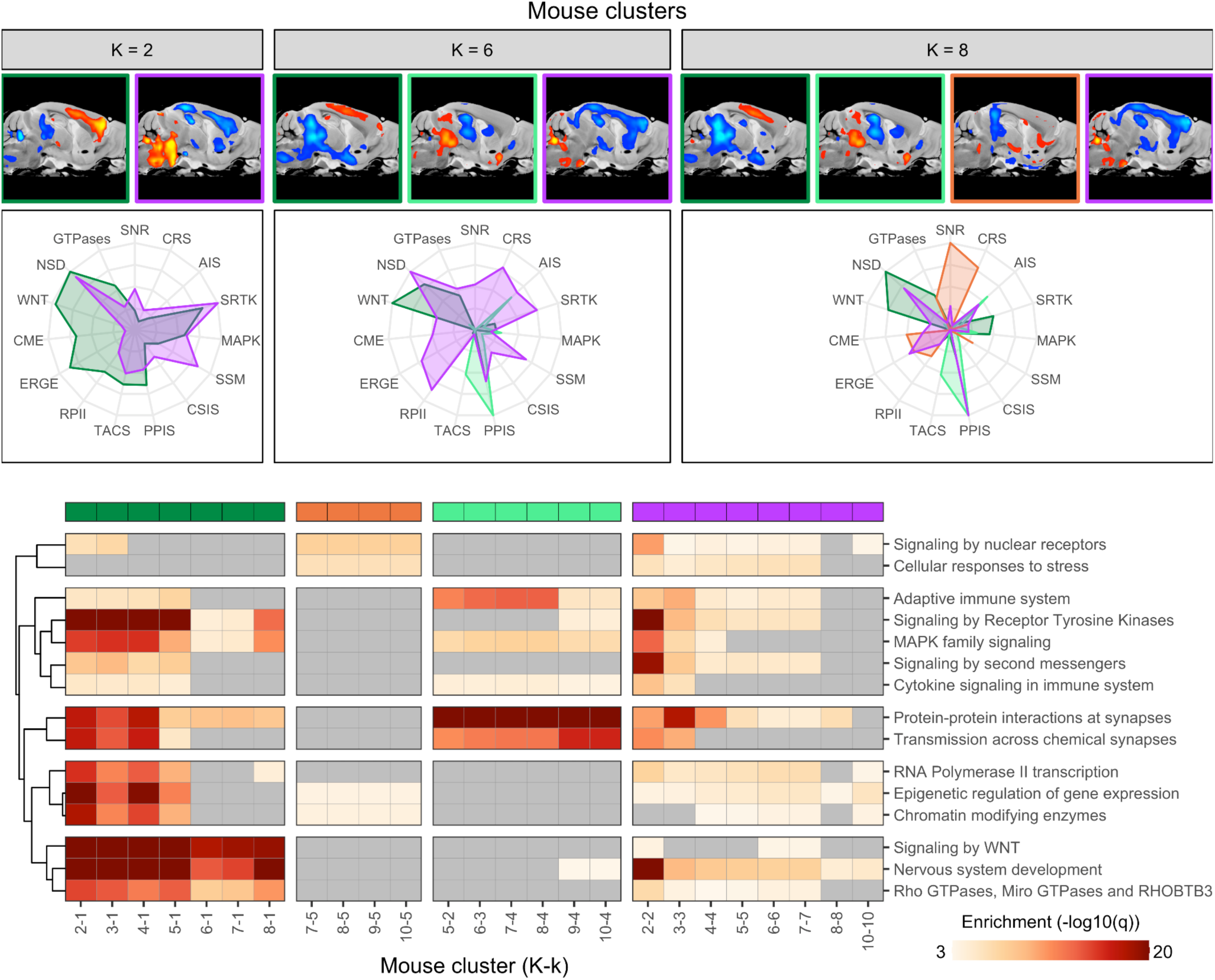
Neuroanatomical clusters are linked to distinct molecular biologies. Mouse neuroanatomical clusters with human matches feature increasingly specific molecular pathway signatures as clustering resolution is increased. (Top) Radar plots of normalized enrichment (-log10(q-value)) per cluster in K = 2, 6, 8 for pathways in the PCA-derived subset. Pathway acronyms and order correspond to the rows in the heatmap. (Bottom) Enrichment heatmap for clusters in stable cross-species subgroups. Clustering resolution (K) increases from left to right within each group. Molecular pathway modules were grouped together based on hierarchical clustering of enrichment patterns across all clusters. Stable clustering groups can be identified as MAPK-WNT (dark green), Androgen Receptor (orange), synaptic-immune (light green), and transcription (purple) based on the molecular programs that emerge at high K.

We subsequently tested whether the matching human clusters in the POND+ sample supported compatible molecular signatures based on the subset of participants with identified rare gene variants. Of the 710 participants in the cohort, genetic information was available for 268 participants and variants were identified for 14 participants. We found that 5-7 variant carriers belonged in clusters matching the mouse MAPK-WNT group, 4 in clusters matching the mouse synaptic-immune group, and 2-3 in clusters matching the mouse transcription group (fig. S7). Using the same enrichment analysis as in the mouse, we evaluated the molecular pathway signatures for these POND+ clusters, and evaluated the correspondence to the matching mouse clusters using an inter-rater agreement strategy. Overall, we find that the correspondence between molecular pathway signatures based on the limited number of variant carriers in our sample is characterized by moderate-to-high specificity with several overlapping pathways between humans and mouse, but low sensitivity with several pathways in the human not showing up in the mouse.

The analyses described above demonstrate that neuroanatomical clusters in both humans and mice are associated with distinct molecular biological programs. We next sought to determine whether these neuro-molecular subtypes were associated with unique diagnostic combinations or clinical and cognitive characteristics. We first examined the diagnostic proportions in the POND+ and HBN clusters and found no difference in diagnostic composition of the clusters (fig. S8). On the mouse side, we evaluated the average SFARI gene score and EAGLE score of the mouse models across the clusters and found that none of the clusters were overly representative of autism based on the genetic information of the composite mouse models (fig. S8). On the human side, we used a battery of behavioural and psychological assessments to evaluate whether the human clusters differed in their clinical and cognitive phenotypes. Starting with a massively univariate approach, we found some nominally significant differences between clusters, but these results did not survive correction for multiple comparisons (fig. S9). We next conducted a multivariate analysis to determine whether clusters could be separated using combinations of behavioural measures. Focusing on the POND+ and HBN clusters with matches in the mouse, we performed separate PCAs in the two cohorts to examine how the clusters separate based on joint behaviour (Fig. 4A). Remarkably, we find evidence that the neuro-molecular clusters distribute similarly based on behaviour in both cohorts, despite a different battery of measures used in each dataset. In particular the cluster means appear to separate along a behavioural axis of IQ and language skills, with the synaptic-immune group exhibiting worse scores. To quantify the behavioural separation of the clusters at the level of individual data points, we then performed pairwise linear discriminant analyses (LDA) to classify the participants into their neuro-molecular groups based on their behavioural scores. The LDA classifiers were able to separate the molecular groups adequately in both the POND+ and HBN cohorts (Fig. 4B). Examining the loadings of the clinical variables onto the axes of maximal separation, we find that the transcription and MAPK-WNT groups are associated with improved performance on measures of IQ, language (OWLS, TOWRE) and adaptive function (ABAS, CGAS, WHODAS) when compared with the synaptic-immune group (Fig. 4C). In the POND+ cohort, the transcription and synaptic-immune groups also feature improved OCD traits (TOCS) compared to the MAPK-WNT group. In the HBN cohort, the synaptic-immune group is also associated with improvements in callous and unemotional traits (ICU) compared to the transcription and MAPK-WNT groups. Together these findings indicate that the neuro-molecular subtypes do not reflect traditional diagnostic categories and are weakly associated with differences in behavioural presentation.

**Fig. 4.**
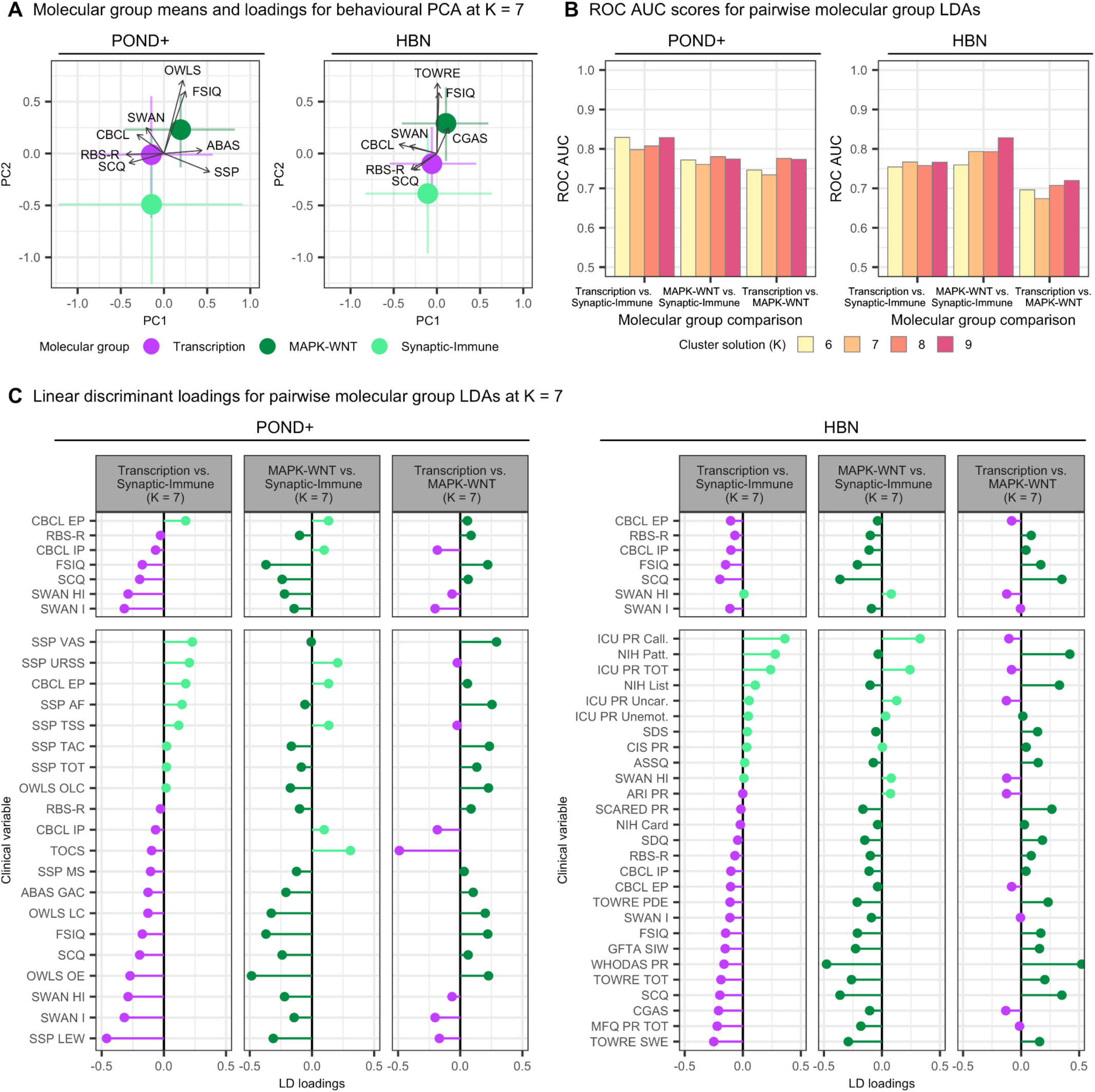
Human neuro-molecular clusters differ along clinical and cognitive axes. (**A**) Molecular group means and loadings for behavioural PCAs in the POND+ and HBN cohorts at K = 7. The first two principal components are shown. Loadings are only displayed for identical or analogous measures in the two datasets, with the exception of the SSP, which is unique to POND. Crosses represent bootstrapped marginal 95% confidence intervals around the cluster means. (**B**) ROC AUC scores for LDA classifiers comparing neuro-molecular groups at K = 6-9 in the POND+ (left) and HBN (right) cohorts. (**C**) Clinical variable loadings associated with the pairwise LDA classification of neuro-molecular clusters at K = 7 in the POND+ (left) and HBN (right) cohorts. The top facets display loadings for the subset of clinical variables available in both POND+ and HBN. The bottom facets display loadings for all clinical variables used in the LDAs. Loadings were calculated using the correlations of the input clinical variables with the learned discriminant axis. All clinical scores were defined such that increasing scores mean improvement along the corresponding domain.

## Discussion

We demonstrate that cross-species structural neuroanatomy identifies conserved signatures of perturbations to molecular pathways that cut across conventional neurodevelopmental diagnoses. Four reproducible neuroanatomical groups emerged, corresponding broadly to MAPK-WNT signalling, androgen receptor signalling, synaptic-immune processes, and transcriptional regulation. Our findings suggest a hierarchical organization in which diverse genetic perturbations converge onto a more limited set of molecular pathways, which in turn produce conserved patterns of brain anatomy that transcend conventional diagnoses.

Our findings are in line with a rich literature from genetics, developmental psychiatry, and transdiagnostic neuroimaging that suggests that conventional neurodevelopmental diagnoses do not map cleanly onto underlying biology(*1*, *9*, *32–36*). Our results extend this work by linking transdiagnostic neuroanatomical organization to molecular pathway signatures through cross-species translation. Moving from diagnoses to more detailed assessments of clinical and cognitive phenotypes revealed some separation amongst our clusters, with the synaptic-immune group exhibiting worse scores along an IQ and language axis. Yet overall phenotypic differences between clusters were more modest than their biological differences, consistent with the well-recognized challenge of mapping brain organization onto behavioural heterogeneity(*37*) and suggesting that molecular convergence is not necessarily accompanied by clinical, behavioural, or cognitive separation.

Our work highlights mouse to human translation. Previous translational studies generally assume that a mouse model either represents a human disorder or it does not. Our results suggest instead that translation is selective: molecularly defined neuroanatomical states exhibit varying degrees of conservation across species. We used MRI as a common phenotype across species, and the expression of common genes(*27*) to provide a shared coordinate space(*38*) in which translation could be quantified. This allowed mechanistic inference to be transferred when supported by the data, rather than assuming universal or absent translation. At higher cluster resolutions, this resulted in approximately 40% of molecularly defined states exhibiting direct cross-species correspondence, suggesting that translational validity is substantial but selective.

We prioritized structural neuroanatomy because it provided a common phenotype that could be measured consistently across hundreds of mouse models and two large human cohorts. Brain anatomy integrates genetic perturbations and environmental effects over development, making it a sensitive tool to capture changes in neural circuitry(*39*). Understanding the cellular bases underlying the mesoscopic brain differences we identified remains important future work, as are further investigations into why these circuits are vulnerable to these pathway perturbations. Additionally, our strategy prioritized breadth of molecular perturbations over depth of phenotyping. Consequently, clustering relied on structural neuroanatomy, with developmental trajectories, behaviour(*9*, *21*), transcriptomics(*40*), and functional imaging(*25*) remaining to be incorporated. Integrating these modalities should refine mechanistic stratification and improve clinical prediction while addressing which modalities are most sensitive to which molecular pathways.

Although additional data modalities will undoubtedly refine these groups, the present framework demonstrates that structural neuroimaging alone contains sufficient information to recover biologically meaningful, cross-species molecular organization. This provides a practical foundation on which richer multimodal stratification can be built. Current precision medicine efforts typically target individual pathogenic variants, limiting recruitment because each variant is rare. By instead stratifying individuals according to convergent molecular pathways, common neuroanatomical signatures may identify substantially larger groups of patients who could benefit from the same therapeutic strategy. Whereas rare pathogenic variants define highly specific genetic disorders and clinical diagnoses summarize heterogeneous behavioural outcomes, the molecular pathway may represent an intermediate level of biological organization shared across both.

Together, these findings suggest that a biologically meaningful unit underlying neurodevelopmental disorders may be neither the diagnostic category nor the causal gene, but convergent molecular pathways reflected in conserved patterns of brain organization. Defining patients according to these pathways offers a route toward mechanism-based clinical trials and precision therapeutics that transcend traditional diagnostic boundaries.

## Acknowledgements

This paper and work are dedicated to the memory of Mark Henkelman whose intellectual insights and contributions were invaluable to this work. Though he is no longer with us, his impact on us and the broader scientific community endures. We greatly miss his presence and are thankful for his mentorship and collaboration on this project and many others.

This work was supported by the Canadian Institute for Health Research and the Ontario Brain Institute (JE,AB,JPL). The Wellcome Centre for Integrative Neuroimaging is supported by core funding from the Wellcome Trust (203139/Z/16/Z and 203139/A/16/Z). For the purpose of open access, the author has applied a CC BY public copyright licence to any Author Accepted Manuscript version arising from this submission.

## Author’s Contributions

JE, AB, YY, MMC, AK, EA, and JPL conceived the study and designed the experiments. JE, AB and YY acquired the data, performed the analyses, the bioinformatics, and statistical comparisons. GAD, JZ, LRQ, SS, MCV, and CL helped with data acquisition and image processing.

MA, JC, SG, AI, JKJ, EK, RS, RW, AK, MJT, EA, and JPL oversaw the recruitment, scanning, and assessment of the human participants.

RA, PS, APAD, LPS, MAB, KQ, NGB, TW, DB, HB, RB, JC, BO, GD, MG, EDB, SE, KDF, SA, JA, KR, JF, JBV, PWF, OP, DHG, RG, SM, JG, CG, LKP DRH, TNH, TLY, YPH, MA, LMI, IL, BK, MB, HJ, HK, HP, JDR, EK, GK, JL, KE, PL, AMB, KM, LSN, BJN, CC, ASN, LMJN, ET, LO, ACA, DP, BB,

HMM, TP, KY, DP, QQX, CMP, AR, DMR, GR, ASS, MWS, LP, CAB, SWS, NAC, SPP, JS, KS, NSK, KJY, GIM, HS, JN, NN, JN, TT, PT, MB, KJ, JV, TK, CLM, JVVW, RW, HB, AWB, and KZ all contributed and/or created mouse models that were prepared and provided for MRI.

JE, AB, and JPL wrote the manuscript with input from all authors. All authors reviewed and approved the final manuscript.

## Supplementary Materials

### Materials and methods

#### Mouse

MRI scanning, registration, and volume measurements were fully automated and unbiased. Experimenters were not blinded to mouse genotypes. All studies were compliant with ethical regulations concerning animal experimentation and approved by the Animal Care Committee at The Centre for Phenogenomics and all the other institutions that provided animals. In total, 3515 different mouse brains were examined from 135 different mouse lines related to autism (table S1). Similar to previous work(*24*), mutant mice from each mouse line were compared with their own wild-type (WT) controls. In all cases the control mice were prepared and provided by the same facility where the mutant mice were created.

Additionally, for some lines we were provided with multiple mutations (i.e. heterozygotes and homozygotes) to compare to WT.

### MRI sequences

#### Perfusion protocol

Perfusions were performed in the lab where the mice we contributed from collaborators around the world (see table S1). All perfusions employed the same protocol, but there were some variations with anesthetic choice depending on the lab. Generally, mice were anesthetized with a ketamine/xylazine mix then intracardially perfused with 30mL of 0.1 M PBS containing 10 U/mL heparin (Sigma) and 2mM ProHance (a Gadolinium contrast agent) followed by 30 mL of 4% paraformaldehyde (PFA) containing 2mM ProHance. After perfusion, mice were decapitated and the skin, lower jaw, ears and cartilaginous nose tip were removed as previously described. The brain within the skull was incubated in 4% PFA containing 2mM ProHance overnight at 4 degrees Celsius then transferred to 0.1M PBS containing 2mM ProHance and 0.02% sodium azide for at least 1 month prior to MRI scanning(*41*, *42*).

#### MRI sequence details

Images were acquired on a 7 Tesla MRI scanner (Agilent, Palo Alto, CA) with either an insert gradient with an inner bore diameter of 6 cm or an outer gradient with a 30 cm inner bore diameter as previously described(*43*, *44*). As data for this project was collected over 10+ years, the anatomical MRI scan that we used evolved over time to increase throughput from 3 mouse brains per scanning session to 16. Additionally, the gradient set and sequence was improved recently to increase the resolution and maintain similar scan times. All sequences gave consistent SNR and tissue contrast with the main difference being the resolution. The original 3 brain sequence resolution was 32 µm isotropic, and the two newer 16 brain sequences had resolutions of 56 µm and 40 µm isotropic. The majority of autism mouse models were investigated with the 56 µm 16 brain sequence.

#### Original sequence – 3 brains per overnight session

Three custom-built solenoid coils were used to image the brains in parallel(*41*, *43*). The sequence used was a T2 weighted Fast Spin Echo (FSE): parameters for this sequence were optimized for high contrast between the gray and white matter. TR – 325 ms, TEs of 10 ms per echo for 6 echoes, with the centre of k-space acquired on the 4^th^ echo, 4 averages, Field-of-view (FOV) of 14 x 14 x 25 mm^3^ and a matrix size of 432 x 432 x 780 yielding an image with 32 µm isotropic voxels. Total imaging time for this MRI sequence was ∼ 12 hrs(*41*, *45*, *46*).

#### Transitional sequence – 16 brains per overnight session

An in-house custom built 16-coil solenoid array was used to acquire anatomical images in parallel, allowing acquisition of the MRI images for 16 samples in one overnight session(*41*, *45*, *46*). Parameters used in this sequence were again optimized for high efficiency and gray/white matter contrast. The sequence was a T2-weighted 3D FSE, with TR=2000 ms, echo train length = 6, TE_eff_=42 ms, FOV of 25 mm × 28 mm × 14 mm, and a matrix size of 450 × 504 × 250. This yielded an isotropic (3D) resolution of 56 µm. In the first phase-encode dimension, consecutive k-space lines were assigned to alternating echoes to move discontinuity related ghosting artifacts to the edges of the FOV. This involved an oversampling in the phase encoding direction by a factor of 2 to avoid the interference of these artifacts. This FOV direction was subsequently cropped to 14 mm after reconstruction. Total imaging time was again ∼12 hours.

#### Current Sequence – 16 brains per overnight session

The same in-house custom built 16-coil solenoid array was used to acquire anatomical images in parallel, again allowing acquisition of the MR images for 16 samples in one overnight session(*41*, *45*, *46*). Parameters used in this improved sequence were again optimized for high efficiency and gray/white matter contrast. The sequence was a T2- weighted, 3-D fast spin-echo sequence was used, with a cylindrical acquisition of k-space(*47*), and with a TR of 350 ms, and TEs of 12 ms per echo for 6 echoes, field-of-view of 20 x 20 x 25 mm^3^ and matrix size = 504 x 504 x 630 giving an image with 40 µm isotropic voxels. Total imaging time for this sequence is currently ∼14 hrs.

### Image registration

After acquisition, we used an image registration pipeline to compare the 3515 brains. The registration was performed in two steps. First, the individual models (or study in the case of multiple models from the same investigator/background) were registered together. Secondly the study averages were then registered to create a global average for the entire sample and bring the data into a common atlas space. The images were aligned using linear (6-parameter fit followed by a 12-parameter fit) and nonlinear registration. Following registration, we resampled the images using the appropriate transform and subsequently created a population average to represent the mean anatomy of the entire study sample. All registrations were performed using a combination of mni_autoreg tools(*48*) and advanced normalization tools (ANTs)(*49*, *50*) using the PydPiper framework(*51*). This process allowed us to deform all images into alignment with each other in an unbiased fashion. It additionally allowed us to analyse the deformations required to take each individual mouse’s anatomy into each step of the pipeline. The Jacobian determinants of the deformation fields representing the registration transforms were used as measures of volume at every voxel.

### Effect size calculation

For computational purposes the data was first resampled to 200um isotropic voxels, which allowed a faster comparison and easier calculations for the large data set used here. For each of the 135 different models, we generated voxel-wise effect size images using both the absolute volume images (mm^3^) and the relative volume images (% of total brain volume). These effect size maps were created comparing the mutant mice in a study to their corresponding wild-type (WT) controls. Effect size was measured in units of standard deviation and was calculated using Glass’s Δ, where the difference between the means is divided by the standard deviation of the WT mice.

### Similarity network fusion and clustering

We used similarity network fusion(*52*) to construct a similarity network of mouse models based on their absolute and relative effect size images. Similarity network fusion was implemented using correlation as the distance metric, with 10 nearest-neighbours, 20 diffusion iterations, and a variance of 0.5 for the local affinity model. We then applied spectral clustering to the fused similarity network to identify clusters of mouse models, from which we extracted solutions from K = 2 to K = 10 clusters. These methods were implemented using the R package ‘SNFtool’(*53*).

### Bioinformatics and enrichment analysis

To evaluate the molecular pathway enrichment of our mouse clusters, we used the genetic information from the models comprising the clusters. Prior to the enrichment analysis, we filtered the set of models for those with precise genetic constructs (116/135). In particular, we removed models that were behavioural (e.g. BTBR and BALB/c), environmental (e.g. VPA and MAR), or based on copy number variations (CNVs), leaving just the single gene models. Given the limited number of single gene models in our clusters, we augmented the gene set associated with each cluster using the STRING database v12.0 (https://string-db.org/)(*29*), which provides information about the protein-protein interaction network for a set of genes. For each cluster, we used the set of single gene models to identify one-hop protein-protein interaction partners. We applied a score threshold of 0.950 to construct the augmented list of genes associated with the given cluster. The score used by the STRING database is an indicator of confidence in the interaction. More information can be found on the STRING database web page or the paper by von Mering et al.(*54*)

Following the augmentation of the cluster gene sets, we evaluated the enrichment of each cluster for 2602 biological and molecular pathway modules from the Reactome Pathway Database (https://reactome.org/)(*31*, *55*). We obtained this set of modules from the Bader Lab at the University of Toronto (http://baderlab.org/, June 2025)(*30*). We calculated the enrichment of each cluster for each module using a hypergeometric test, implemented with the ‘tmod’ package in the R programming language. For the background set of genes required by these statistical tests, we used a set of 17,970 genes from the sagittal data set of the Allen Mouse Brain Atlas. For each cluster, we corrected the p-values for multiple comparisons across the 2,602 molecular pathway modules using the Benjamini-Hochberg method for controlling the false discovery rate(*56*). The resulting q-values were transformed using a –log10 transformation. To create a subset of pathways for analysis, we first filtered for pathways belonging to the first level of the Reactome pathway hierarchies. We then applied a PCA using the enrichment of this initial subset of pathways across all 54 mouse clusters to determine which pathways were the most enriched in our dataset. The final set of 15 pathways used in our analyses were obtained by filtering for those pathways with an absolute score of 5 or more along the first principal component (table S3).

### Human

#### Participants

As our primary cohort we used a joint dataset from the Province of Ontario Neurodevelopmental Disorders (POND) Network (n = 750) and an imaging cohort from Dr. Margot Taylor’s group at the Hospital for Sick Children (n = 480). The dataset included 710 (57.7%) participants diagnosed with a neurodevelopmental disorder and 520 (42.3%) participants who were neurotypical. In the NDD cohort, the primary diagnoses were split between autism (n = 399), ADHD (n = 228), OCD (n = 73), anxiety (n = 2), intellectual disability (n = 1), Tourette syndrome (n = 2), fragile X syndrome (n = 1), and other (n = 4). For the purposes of our analyses, we grouped the participants who did not have a diagnosis of autism, ADHD, or OCD into a general category named “Other”. The participants were between the ages of 2 and 65 years (mean age: 13.6 +/- 7.0) with 802 (65.2%) males and 428 (34.8%) females. All participants and their parents were able to complete the testing protocols in English and had no contraindications to magnetic resonance imaging (MRI). Participants in the clinical groups met the DSM criteria for their respective diagnosis and diagnoses were supported by gold-standard assessments [autism: Autism Diagnostic Observation Schedule–2 (ADOS)(*57*) and Autism Diagnostic Interview–Revised (ADI-R)(*20*); ADHD: Parent Interview for Child Symptoms (PICS)(*58*)]. Participants in the neurotypical group did not have a neurodevelopmental, psychiatric and/or neurological diagnosis or any first-degree relatives with a neurodevelopmental condition and were born after 35 weeks’ gestation. Informed consent was provided by participants (when they had the capacity to consent) or their guardians, and assent was obtained from all participants as per institutional ethics board guidelines. Institutional research ethics boards approvals were received for the study.

As a replication dataset for our human findings, we used the Child Mind Institute Healthy Brain Network (HBN) cohort, which consisted of 1,015 participants, with 873 (86.0%) NDD participants and 142 (14.0%) controls. The NDD group contained 105 participants with autism, 726 with ADHD, 12 with OCD, 18 with intellectual disability, and 12 with Tourette syndrome. Ages ranged from 5 to 21 years, with a mean of 10.3 +/- 3.4. The cohort contained 724 (71.3%) males and 291 (28.7%) females.

#### Imaging data

We used structural MRI to obtain T1-weighted (T1w) images of whole-brain anatomy. In the POND+ cohort, the images were collected at three separate sites across Ontario: The Hospital for Sick Children (n = 1072) and Holland Bloorview Kids Rehabilitation Hospital (n = 19) in Toronto, and Queen’s University in Kingston (n = 143). Most of the scans acquired at the Hospital for Sick Children were obtained using a 3-Tesla Siemens Trio TIM (n = 711), which was later upgraded to a Siemens Prisma Fit scanner (n = 361). Likewise, most of the images from Queen’s University were also obtained using a Siemens Trio TIM scanner (n = 107) before an upgrade to a Siemens Prisma Fit (n = 36). All images acquired at Holland Bloorview were obtained using a Siemens Prisma Fit (n = 19).

In the HBN cohort, the images were collected at three sites in New York: the Rutgers University Brain Imaging Center (n = 539), the CitiGroup Cornell Brain Imaging Center (n = 443), and the City College of New York (n = 33). Images from Rutgers University were obtained using a 3-Tesla Siemens Trio TIM scanner, while the others were obtained using a 3-Tesla Siemens Prisma Fit.

#### Image registration

We used an image processing pipeline for image registration and deformation-based morphometry to derive local measures of absolute and relative volume differences for each participant. Prior to image registration, the T1w images of all participants in both the POND+ and HBN datasets were pre-processed using a pipeline for iterative N3 bias field correction (https://github.com/coBrALab/iterativeN3 and then manually assessed for motion artifacts using the QC procedure described in Bedford et al 2020(*59*). Only those images that successfully passed both stages were selected for image registration and subsequent analysis. Model building for image registration was then performed with a model schedule of rigid[1], similarity[1], affine[1], non-linear[4], and a gradient step of 0.25 (https://github.com/CoBrALab/optimized_antsMultivariateTemplateConstruction). The starting target for the registration was taken to be the MNI ICBM non-linear 09c 1mm model, which was extracted, re-cropped, and upsampled to an isotropic resolution of 0.5mm. Following image registration, post-processing of the resulting deformation fields produced absolute Jacobian determinant images encoding the contributions of the affine transform between the participants and the template, and relative Jacobian determinant images with contributions of residual affine components removed. The absolute and relative Jacobian images were then smoothed using a Gaussian kernel with a full-width at half-maximum of 2 mm.

#### Calculating effect sizes

We generated effect size images for each participant in the NDD group using both absolute and relative Jacobian determinant images separately. We began by controlling for the effects of imaging site and scanner by regressing the Jacobian determinants against these terms using a simple linear model on a voxel-wise basis and extracting the model residuals. Then, using the Jacobian images from the typically developing control participants, we generated a normative growth model at every voxel. Our model included a covariate for sex as well as a natural cubic spline with three degrees of freedom for age. We then generated the absolute and relative effect size image for the NDD participants by normalizing their Jacobian determinant images against the voxel-wise normative models.

#### Similarity network fusion and clustering

We applied similarity network fusion and spectral clustering to the absolute and relative human effect size images to identify clusters of human participants, as described previously for the mouse models. Parameters were identical for the mouse and human.

#### Bioinformatics and enrichment analysis

To perform a molecular pathway enrichment analysis on the POND+ human clusters, we used those participants in our cohort with identified genetic variants. Of the 710 NDD participants in our study whose MRI images survived QC, 268 participants had sequencing data(*14*). Of these, only 18 had identified rare autism associated genetic variants. For the enrichment analysis, we further reduced this set down to 14 participants who had single gene variants. We performed the enrichment analysis in the same way that we did for the mouse clusters, i.e. first by augmenting the cluster gene sets using the STRING database, and then running hypergeometric tests for the pathway modules from the Reactome Pathway Database.

#### Behavioural analysis

For the POND human cohort, we used a battery of behavioural assessments consisting of: the Adaptive Behaviour Assessment System (ABAS-II) (5-21 years) General Adaptive Composite (GAC) score; the Bruininks-Oseretsky Test (BOT-2) balance, body coordination, and bilateral coordination standard scores; the Child Behavior Checklist (CBCL) internalizing and externalizing problems T-scores; the Developmental Neuropsychological Assessment (NEPSY-II) 5-16 years, Affect Recognition (AR), Memory for Faces (MF), Memory for Faces Delayed (MFD), Theory of Mind (TM) scaled scores, MF vs. MFD contrast scaled score; the (Oral and Written Language Scales) OWLS-II Listening Comprehension (LC), Oral Expression (OE), and Oral Comprehension (OC) standard scores; the Repetitive Behaviour Scales - Revised (RBS-R) overall score; the Social Communication Questionnaire (SCQ) lifetime total score; the Short Sensory Profile 1 tactile sensitivity raw score, taste/smell sensitivity raw score, movement sensitivity raw score, under responsive/seek sensation raw score, auditory filtering raw score, low energy/weak raw score, visual/audio sensitivity raw score, and total raw score; the (Strength and Weaknesses of ADHD Symptoms and Normal Behaviours (SWAN) Rating Scale; the Toronto Obsessive-Compulsive Scale (TOCS); and a composite measure of IQ based on the Wechsler Abbreviated Scale of Intelligence (WASI-II), the Wechsler Intelligence Scale for Children (WISC), and the Stanford-Binet.

For the HBN cohort, we used a battery of assessments with high completion rates. These included: the Children’s Global Assessment Scale (CGAS); the Autism Spectrum Screening Questionnaire (ASSQ); the Affective Reactivity Index (ARI) Parent Report total score; the Screen for Child Anxiety Related Disorders (SCARED) Parent Report total score; the Mood and Feelings Questionnaire (MFQ) Parent Report total score; the Strengths and Difficulties Questionnaire (SDQ) total difficulties score; the WHO Disability Assessment Schedule (WHODAS) Parent Report score; the Columbia Impairment Scale (CIS) Parent Report total score; the Sleep Disturbance Scale (SDS) total T-score; the Inventory of Callous-Unemotional Traits (ICU) Parent Report callousness, uncaring, unemotional subscale scores and total score; the NIH Toolbox dimensional change card sort, flanker inhibitory control/attention, pattern comparison process speed, and list sorting working memory scaled scores; the Test of Word Reading Efficiency (TOWRE-2) phonemic decoding efficiency, sight word efficiency, and total word reading efficiency scaled scores; the Goldman-Fristoe Test of Articulation (GFTA) sounds-in-words standard score; the Youth Self Report (YSR) total T-score; and the same scores as POND for the SCQ, RBS-R, SWAN, CBCL, and IQ.

In each cohort, we first evaluated the distributions of these measures across cluster solutions using a massively univariate implementation of Kruskal-Wallis tests: for each of the behavioural scales and each cluster solution in K = 2-10, a Kruskal-Wallis test was used to evaluate the score distributions across clusters. For each cluster solution the p-values were corrected for multiple comparisons across behavioural measures using the false discovery rate method.

We then employed a multivariate approach, deploying a set of pairwise linear discriminant analyses (LDA) to compare clusters with matches to mouse molecular cluster groups. We selected a subset of behavioural assessments to use as input features based on participant completion of the measures. For POND+, the 20 variables with the highest completion rates were selected, resulting in 200 participants with complete data. For HBN, the top 27 variables were selected, resulting in 581 participants with complete data. For each K and each pairwise comparison, the performance of the model was quantified using the area under the curve (AUC) of the associated receiver operating characteristic (ROC) curve.

### Linking mouse and human clusters

#### Evaluating cluster similarity in gene expression space

We evaluated the similarity between mouse and human clusters by first computing absolute and relative centroid images for the clusters, and then decomposing these centroids into a common space built from mouse-human homologous genes. This common space was constructed using the openly accessible Allen Human Brain Atlas(*60*) and Allen Mouse Brain Atlas datasets(*61*) from the Allen Institute for Brain Science, as described in Beauchamp et al., 2022(*27*). Here we generated a total of 100 gene expression latent spaces using this method. Each space consisted of 200 latent variables and each of these variables was represented as an image in an appropriate imaging space. The mouse latent variable images existed in the native Allen Common Coordinate Framework v3 (CCFv3)(*62*). For the human latent variables derived from the AHBA microarray data, we mapped each microarray sample to a voxel in MNI space using a set of coordinates carefully registered to MNI ICBM nonlinear symmetric 09c space (https://github.com/gdevenyi/AllenHumanGeneMNI).

For each of the mouse and human clusters, we first created a set of absolute and relative centroid images by evaluating the voxel-wise mean of the absolute and relative effect size images of those mouse models or human participants in the cluster. These images were transformed and resampled to the appropriate imaging spaces described in the previous paragraph. For each of these centroids, we then created a mask identifying which of the voxels would contribute to the gene space decomposition. These masks were constructed to filter for the top 20% of voxels based on the absolute value of the centroid effect sizes. While we used the absolute values to determine which voxels should contribute to the mask, we kept track of the sign of the contributing voxels by encoding the mask using −1 for negative voxels and +1 for positive voxels. These masks were generated separately for the absolute and relative Jacobian centroid images. We then used these signed masks to construct gene expression signatures for the mouse and human clusters: For each cluster and each class of Jacobians, we separated the mask into its positive and negative components. In cases where only positive or negative values were present, only that sign was used. We then applied the positive and negative masks to each of the 200 latent space variable images to select for those voxels “belonging” to the cluster. The values of each latent space variable were averaged within these masks to create a positive and a negative gene expression signature for the cluster. Subsequently, we evaluated the pairwise similarity between all mouse and human clusters using the Pearson correlation coefficient separately for the positive and negative gene expression signatures, resulting in positive and negative similarity matrices. These were then averaged to create the similarity matrix for the given class of Jacobians and the given latent space. We repeated this process for each of the 100 latent spaces, resulting in 100 similarity matrices based on absolute Jacobians and 100 similarity matrices based on relative Jacobians. We obtained a final similarity matrix by first computing the average across all latent spaces for each Jacobian type, and then taking the average across Jacobians.

#### Evaluating the significance of cluster similarity

To evaluate the significance of the correlation between pairs of clusters (mouse vs. human and human vs. human), we implemented a permutation-based approach. In each permutation, we used sampling without replacement to shuffle the association between the human participants and their cluster assignment for each of the cluster solutions from K = 2 to K = 10. In the case of mouse vs. human comparisons, the cluster assignments of the mouse models being compared were held constant. In the case of POND+ vs. HBN comparisons, the assignments of the POND+ participants were shuffled while the cluster assignments of the HBN participants were held constant. We then repeated the process described in the previous section to evaluate the correlation between the mouse (or human HBN) clusters and the permuted POND+ human clusters. However, for each of the K-cluster solutions in either species, we only evaluated the permuted similarity to the (K-1), K, and (K+1) cluster solutions in the other species. This procedure was repeated for 500 permutations. To evaluate the statistical significance of mouse-human cluster correlations, we constructed a separate empirical null distribution for each pair of cluster solutions. For instance, to evaluate the significance of pairwise correlations between the clusters in the human J-cluster solution and those in the mouse K-cluster solution, we used all J-by-K correlations across all 500 permutations to generate the null distribution. Using these empirical null distributions, we calculated a p-value for the true cluster correlations by evaluating the associated tail probability.

## Data and Code Availability

For the Mouse Data - Currently 27 of the models are available on brain code https://www.braincode.ca/. On publication of this work the remaining mouse models will be uploaded and available on that platform as well. For the Human Data - The primary cohort can be found through the POND network on Braincode https://www.braincode.ca/ and is consistently being updated. The Replication Cohort can be found at the Child Mind Institute Healthy Brain Network (HBN) https://healthybrainnetwork.org/. Mouse and Human MRI analysis was done in R (https://www.r-project.org) using the RMINC toolkit (https://github.com/Mouse-Imaging-Centre/RMINC). Prior to image registration, the T1w images of all participants in both the POND and HBN datasets were pre-processed using a pipeline for iterative N3 bias field correction (https://github.com/coBrALab/iterativeN3 and then manually assessed for motion artifacts using the QC procedure described in Bedford et al 2020^24^. The code that was used for the clustering analysis can be found in GitHub - https://github.com/abeaucha/ClusteringAutism/

## Supplementary Figures

**Fig. S1.**
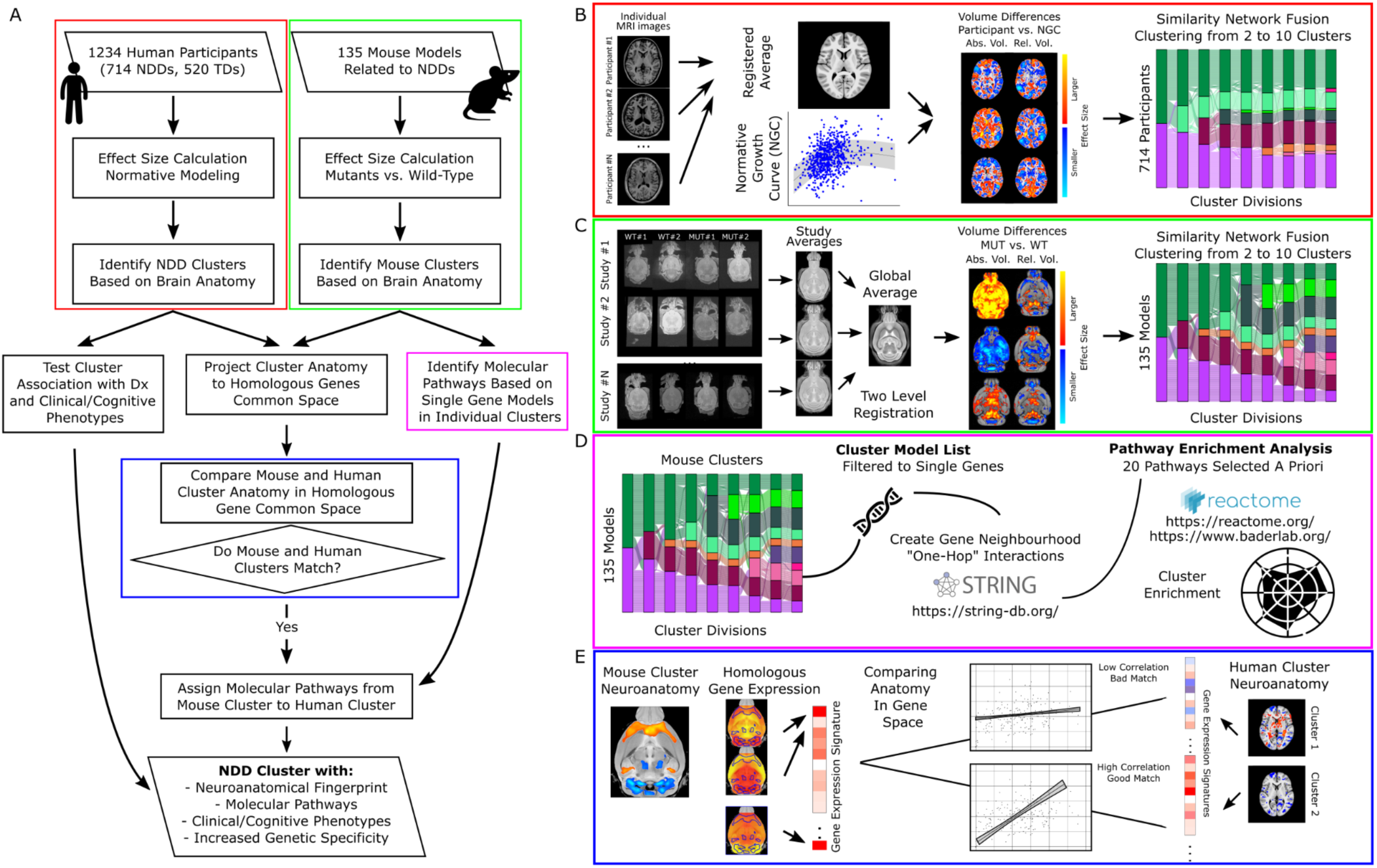
Detailed overview of project workflow. **(A)** On both the human and mouse sides of the analysis, we implemented an image processing workflow in which we used image registration to align the individual brain images to a species-specific common consensus average space, generated effect-size images for the human participants and mouse models, and separately identified clusters of human participants and mouse models based on neuroanatomical similarity. **(B)** For the human data, all images were registered together in a single stage. To compute the effect size images for the NDD participants, we first generated voxel-wise normative growth curves using the subset of typically developing controls. We then normalized each NDD participant against these growth curves. Clusters were identified using similarity network fusion and spectral clustering. **(C)** For the mouse data, image registration was done in two stages, by first generating study-specific consensus average spaces, and then subsequently registering those spaces to a global average. We generated effect size images for each study by normalizing the mutant mice against their respective wild-type controls. Clustering was performed the same as for humans. **(D)** Mouse clusters were linked to molecular pathways using a combination of the STRING database (https://string-db.org)^15,16^, publicly available gene lists the Bader group (https://www.baderlab.org)^19^, and the Reactome database (https://reactome.org)^17,18^. Further details of the parameters used to assign these gene lists and pathways can be found in the methods. **(E)** We evaluated the degree of similarity between the mouse and human clusters using freely available spatial gene expression data sets from the Allen Human Brain Atlas and Allen Mouse Brain Atlas. We first created a centroid image for each mouse and human cluster, which we then decomposed into a set of mouse-human orthologous genes, resulting in a gene expression signature for each cluster. We then quantified the similarity between a mouse cluster and a human cluster by correlating their respective gene expression signatures. In doing so, we can link certain mouse and human clusters based on the similarity of their neuroanatomical morphology, and thus assign specific genetics and molecular pathway information directly to human participants.

**Fig. S2.**
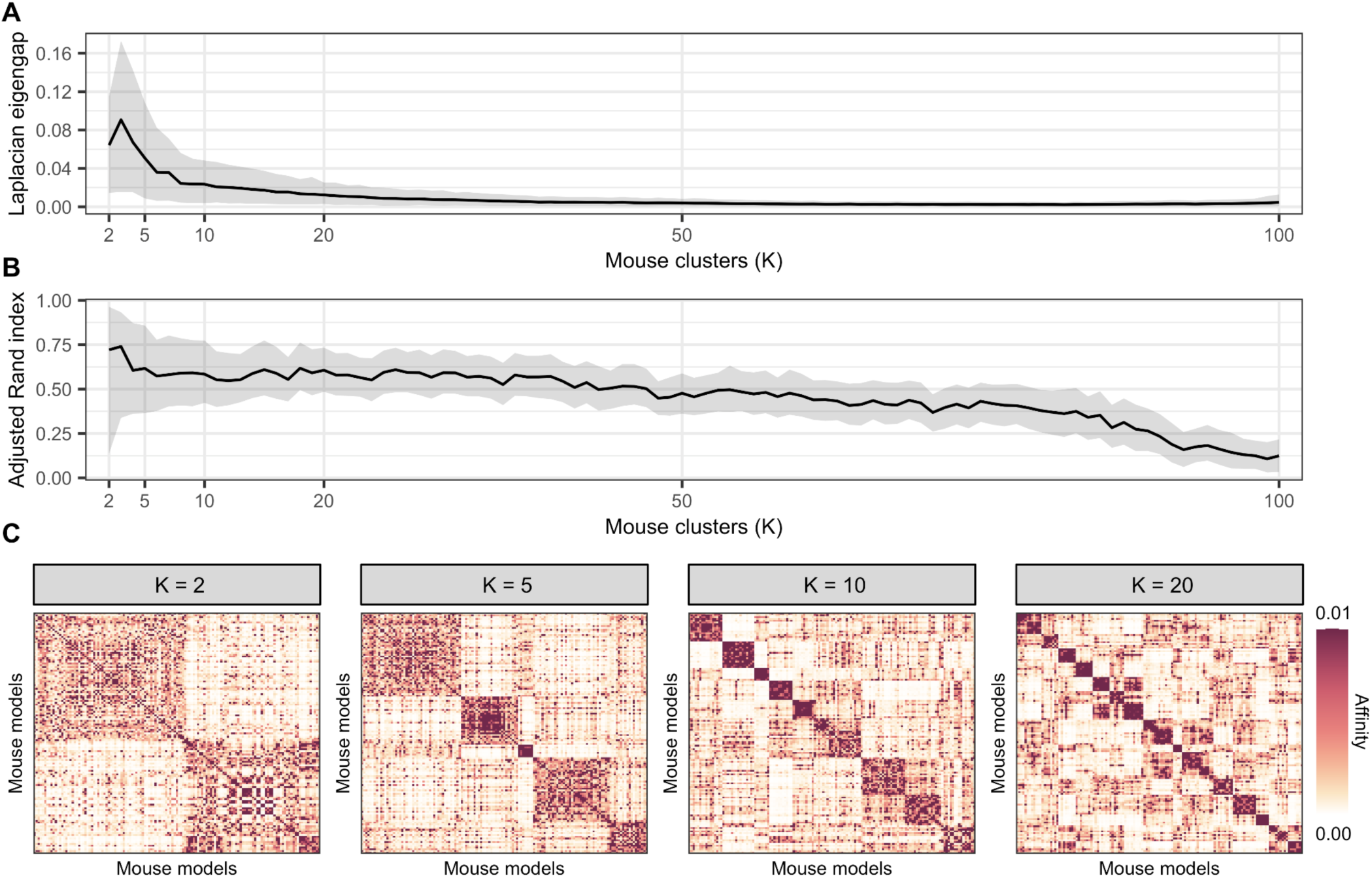
Mouse clusters exhibit multiscale organization. (A) Distribution of the graph Laplacian eigengap across mouse clustering solutions with K = 2-100. Parameter variance was estimated using repeated subsampling (B = 500) of 80% of mouse models (n = 108) followed by similarity network construction and clustering. At low K, cluster solutions feature higher separability on average but separability is sensitive to the subset of mouse models included. As K increases from K = 2 to K = 20, the clusters gradually become less separable on average, but robustness to mouse model inclusion increases. At higher K, clusters cease to be separable regardless of which mouse models are included in the analysis. (B) Distribution of the adjusted Rand index (ARI) across mouse clustering solutions. The ARI was calculated for each K by repeatedly (B = 500) sampling 80% of mouse models and comparing each sample against the true partition obtained by clustering the full set of models. The distribution of ARI suggests moderate cluster stability with respect to mouse model inclusion from K = 2 to around K = 50, with high sensitivity to model inclusion at low K. (C) Mouse model affinity matrices for clustering solutions at K = 2, 5, 10, and 20 highlighting well-defined clustering structure over this range of scales.

**Fig. S3.**
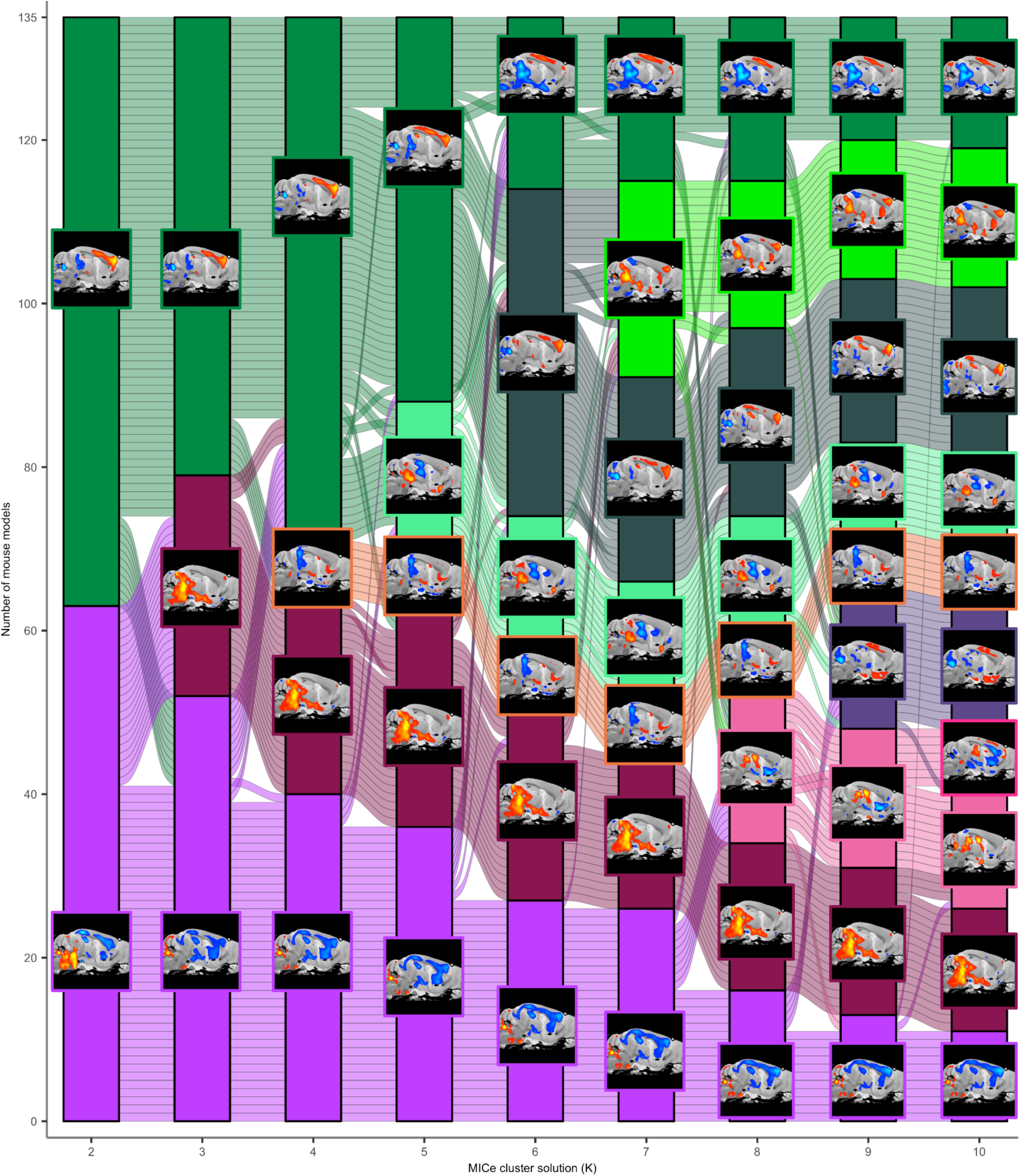
Mouse clusters and neuroanatomy from K = 2 to K = 10. Sankey diagram of mouse model clusters for K = 2-10. Sagittal brain slices indicate the relative effect size centroid for each cluster.

**Fig. S4.**
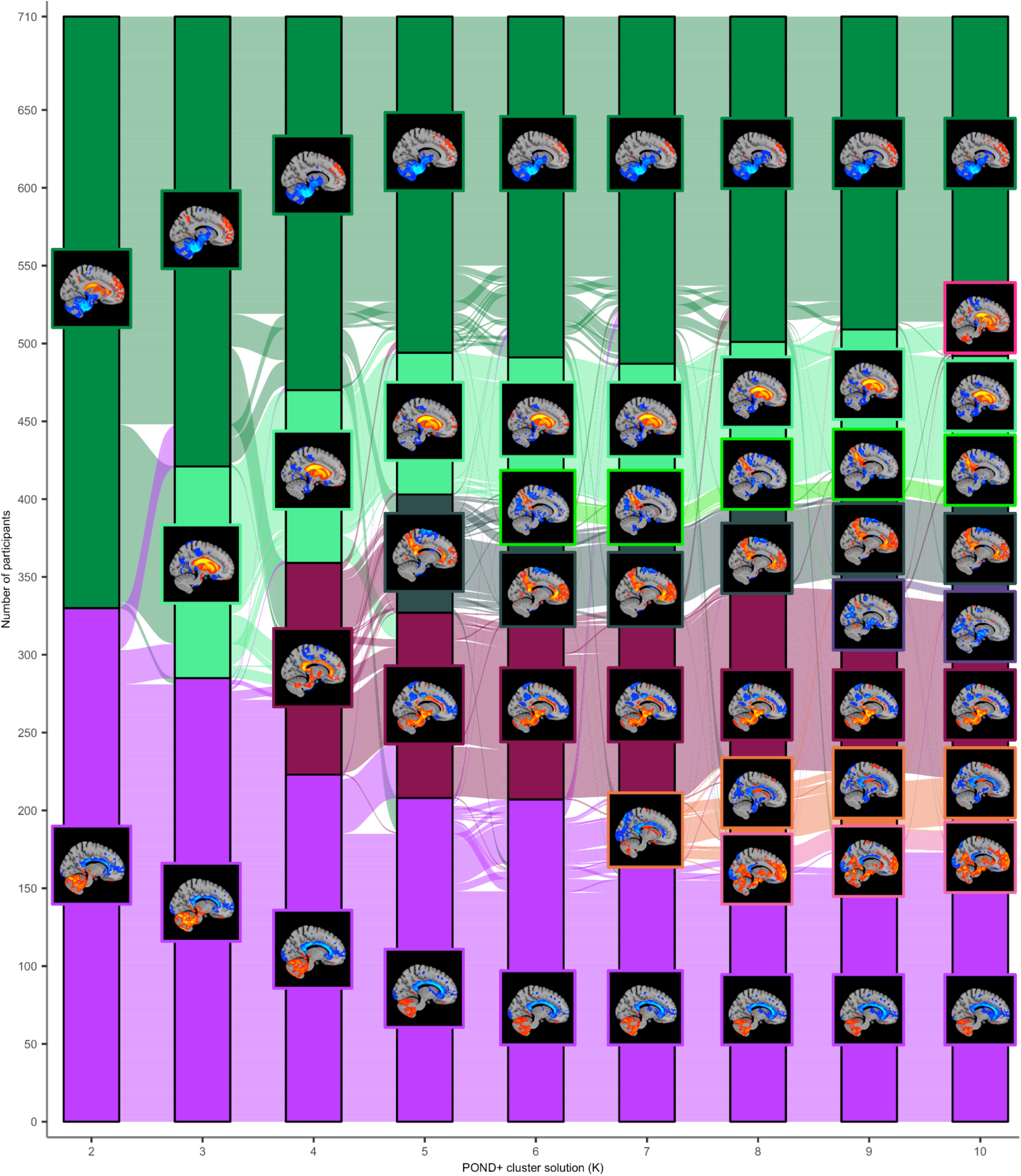
POND+ clusters and neuroanatomy from K = 2 to K = 10. Sankey diagram of POND+ clusters for K = 2-10. Sagittal brain slices indicate the relative effect size centroid for each cluster. Where possible, the colour palette reflects the mouse clusters with the highest similarity.

**Fig. S5.**
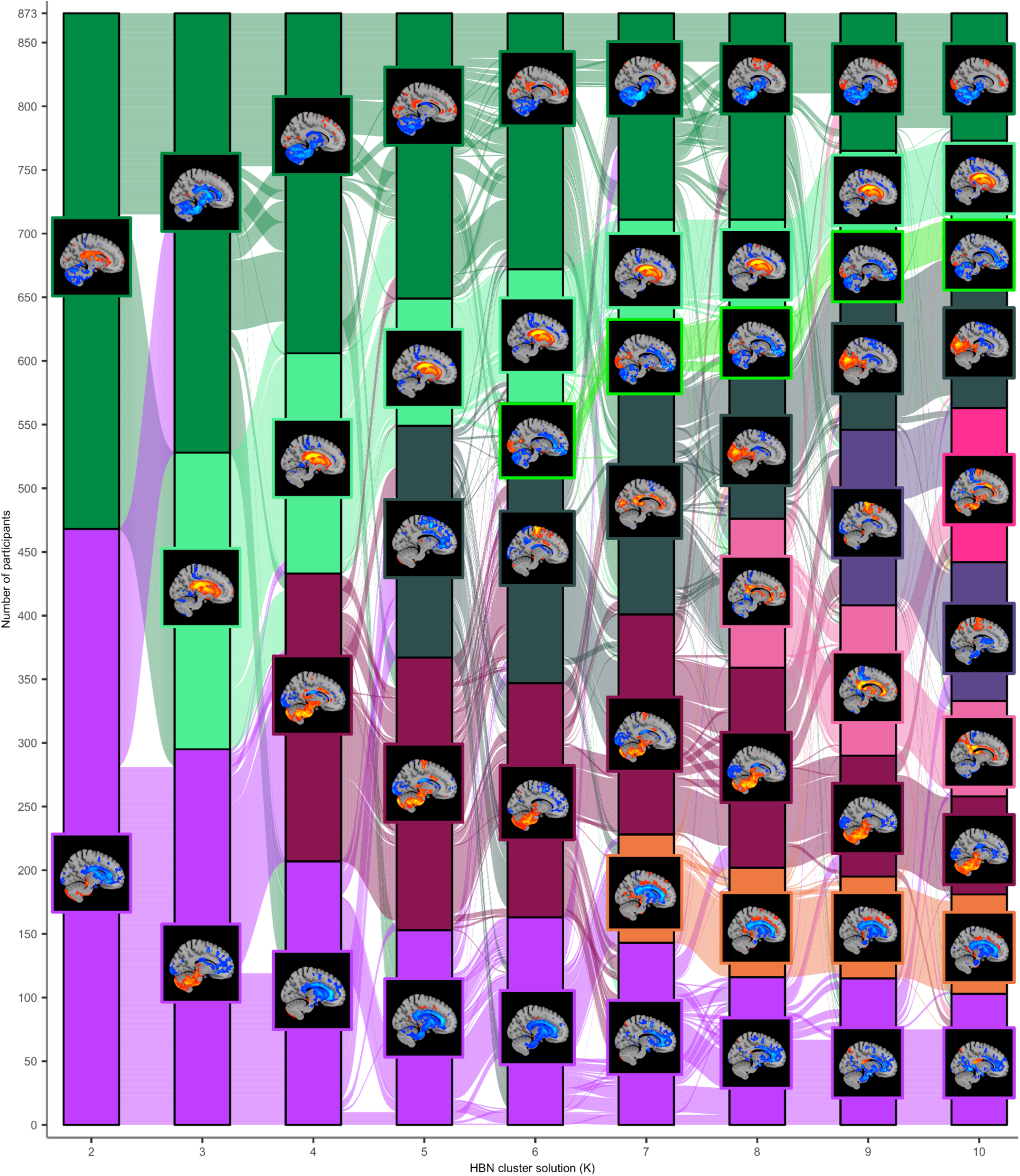
HBN clusters and neuroanatomy from K = 2 to K = 10. Sankey diagram of HBN clusters for K = 2-10. Sagittal brain slices indicate the relative effect size centroid for each cluster. Where possible, the colour palette reflects the POND+ clusters with the highest similarity.

**Fig. S6.**
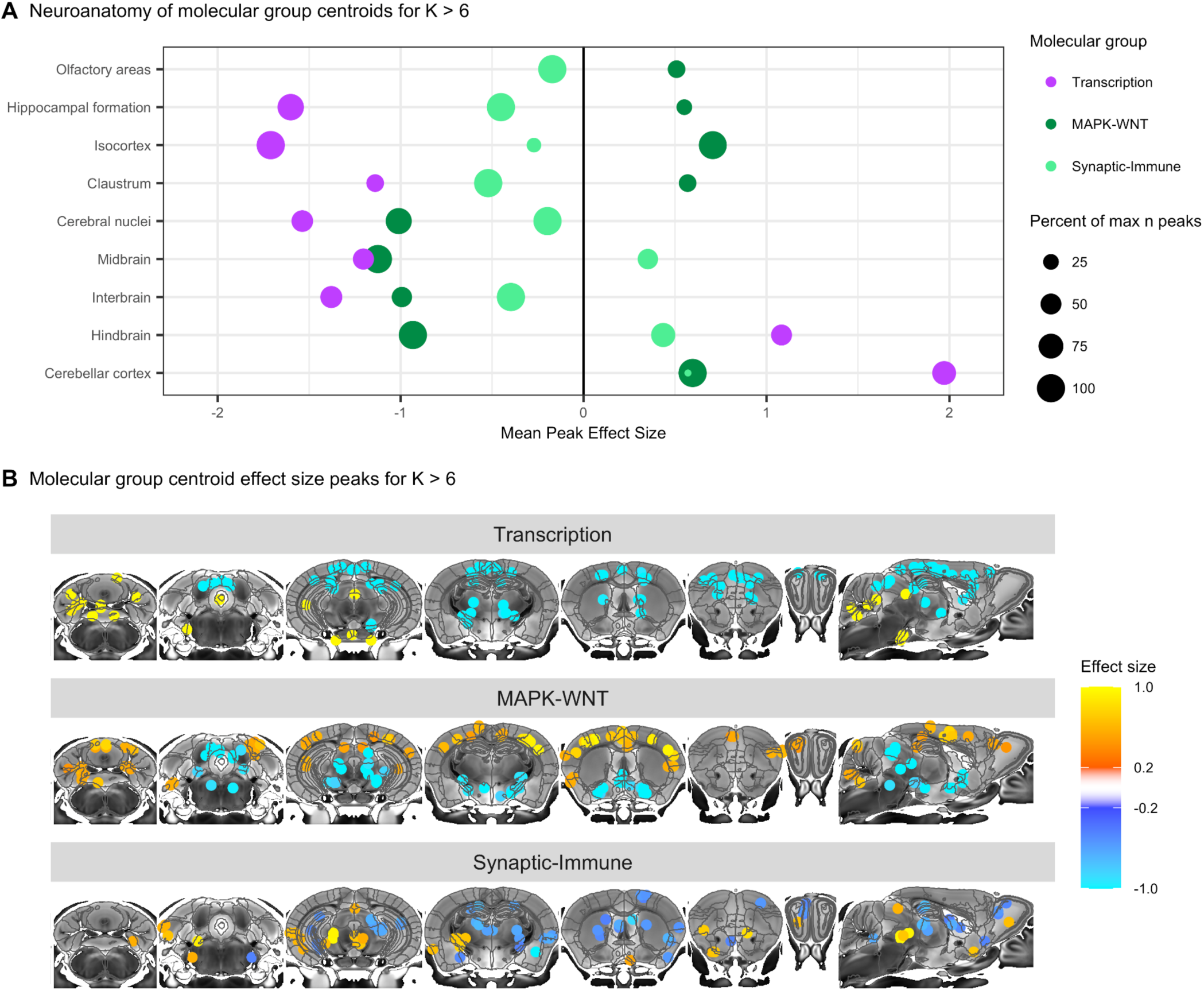
Neuroanatomical patterns associated with mouse molecular pathway groups. (**A**) Distributions of mean centroid peaks across broad neuroanatomical regions for mouse clusters with K > 6 in the transcription, MAPK-WNT, and synaptic-immune molecular cluster groups. Point sizes indicate the percentage of the maximum number of peaks found in each brain region. (**B**) Neuroanatomical slices displaying peak centroid values for mouse clusters with K > 6 in the molecular cluster groups. The right-most slice is a midline sagittal slice.

**Fig. S7.**
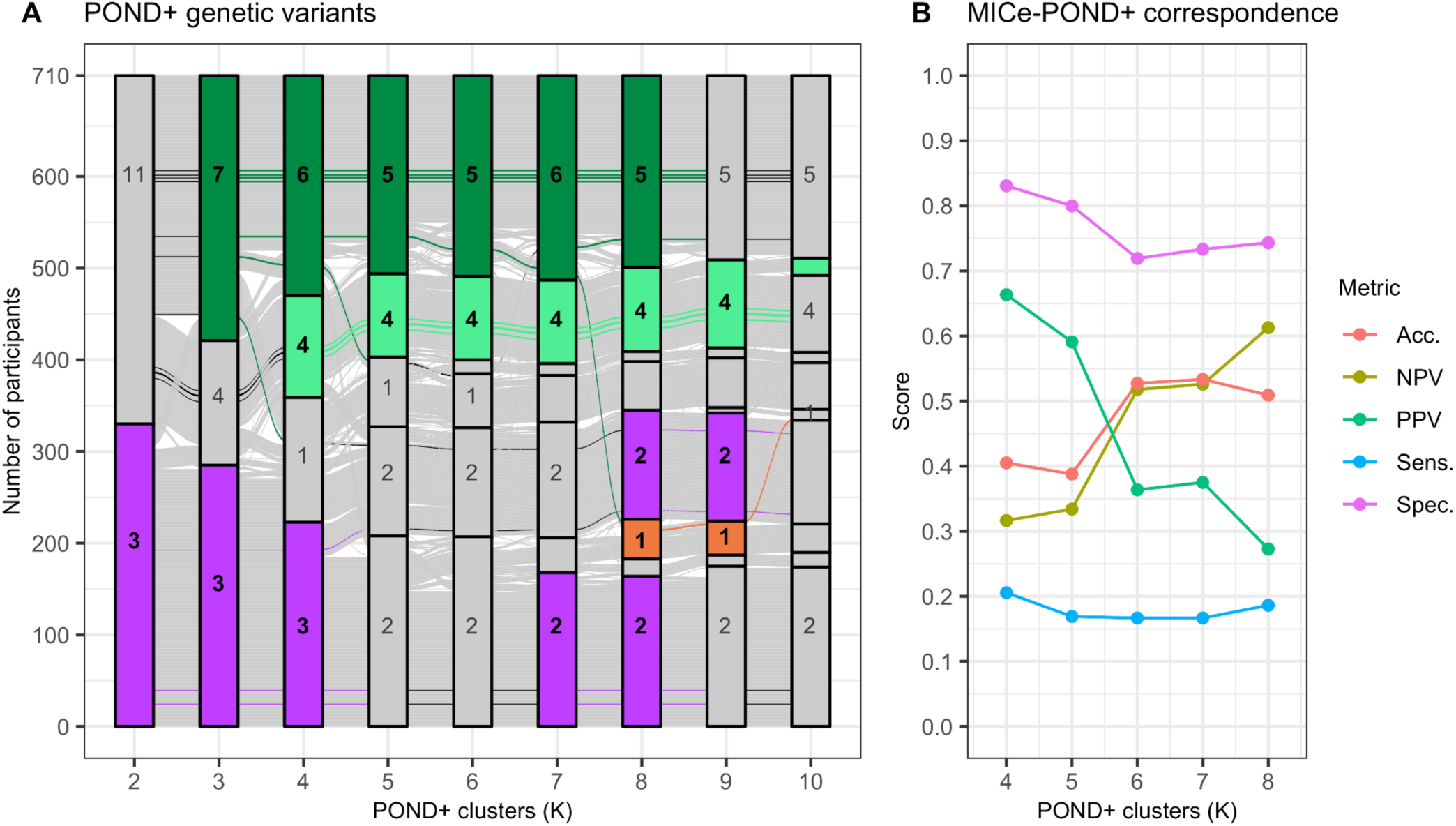
POND+ genetic variant carriers reflect moderate molecular pathway agreement with mouse clusters. (**A**) POND+ clusters with identified variant carriers per cluster (numbered) and matching mouse cluster groups (coloured). Grey clusters have no significant mouse match. Coloured alluvials correspond to individual variant carriers. (**B**) Correspondence analysis between the human and mouse pathways for the clusters with cross-species matches. Correspondence was evaluated at each relevant K using a confusion matrix for the MAPK-WNT, synaptic-immune, and transcription groups by treating the enriched mouse pathways as ground truth and the enriched human pathways as predictions. Each confusion matrix was used to compute the accuracy (Acc.), negative predictive value (NPV), positive predictive value (PPV), sensitivity (Sens.) and specificity (Spec.). At each K, these metrics were then averaged across the three molecular cluster groups (MAPK-WNT, synaptic-immune, transcription) to generate a final overall measure of agreement.

**Fig. S8.**
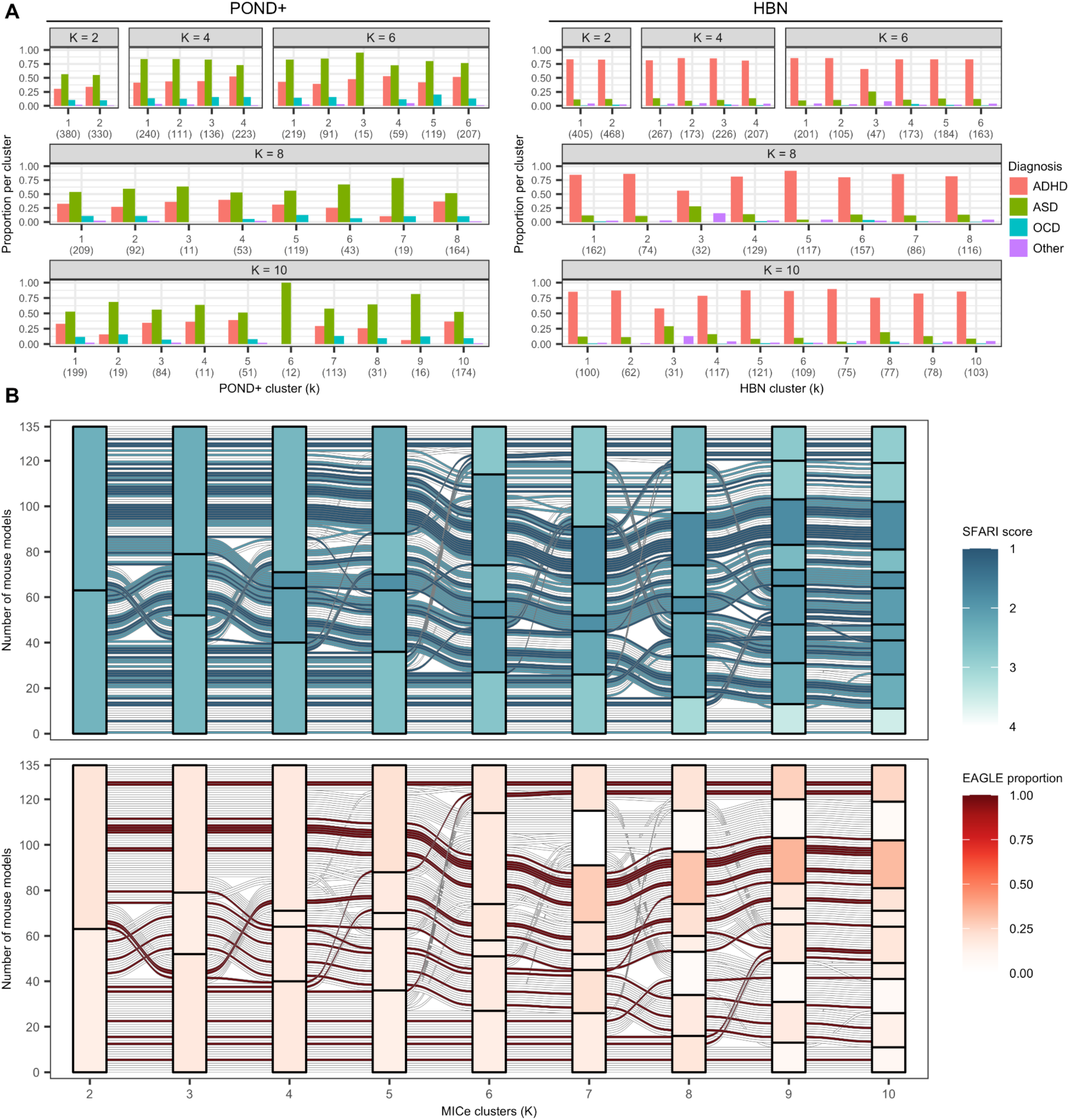
Neuro-molecular clusters do not separate into traditional diagnoses. (**A**) Diagnostic proportions per cluster for K = 2, 4, 6, 8, 10 in the POND+ (left) and HBN (right) cohorts. Cluster sizes are indicated in parentheses along the x-axes. Clusters do not cleanly separate traditional diagnostic groups, with the exception of small clusters at higher K (e.g. POND+ cluster 10-6). (**B**) Mouse SFARI (top) and EAGLE (bottom) scores for mouse models and clusters across K = 2-10. SFARI scores for the individual mouse models are displayed using the alluvials and mean SFARI scores are presented for the clusters. The EAGLE scores for individual mouse models were binarized using a threshold of 11, with scores greater than 11 indicating a strong link to autism (red alluvials). The proportion of mouse models with suprathreshold scores is displayed for each of the clusters. None of the clusters is disproportionately associated with autism specifically, based on genes of the composite mouse models.

**Fig. S9.**
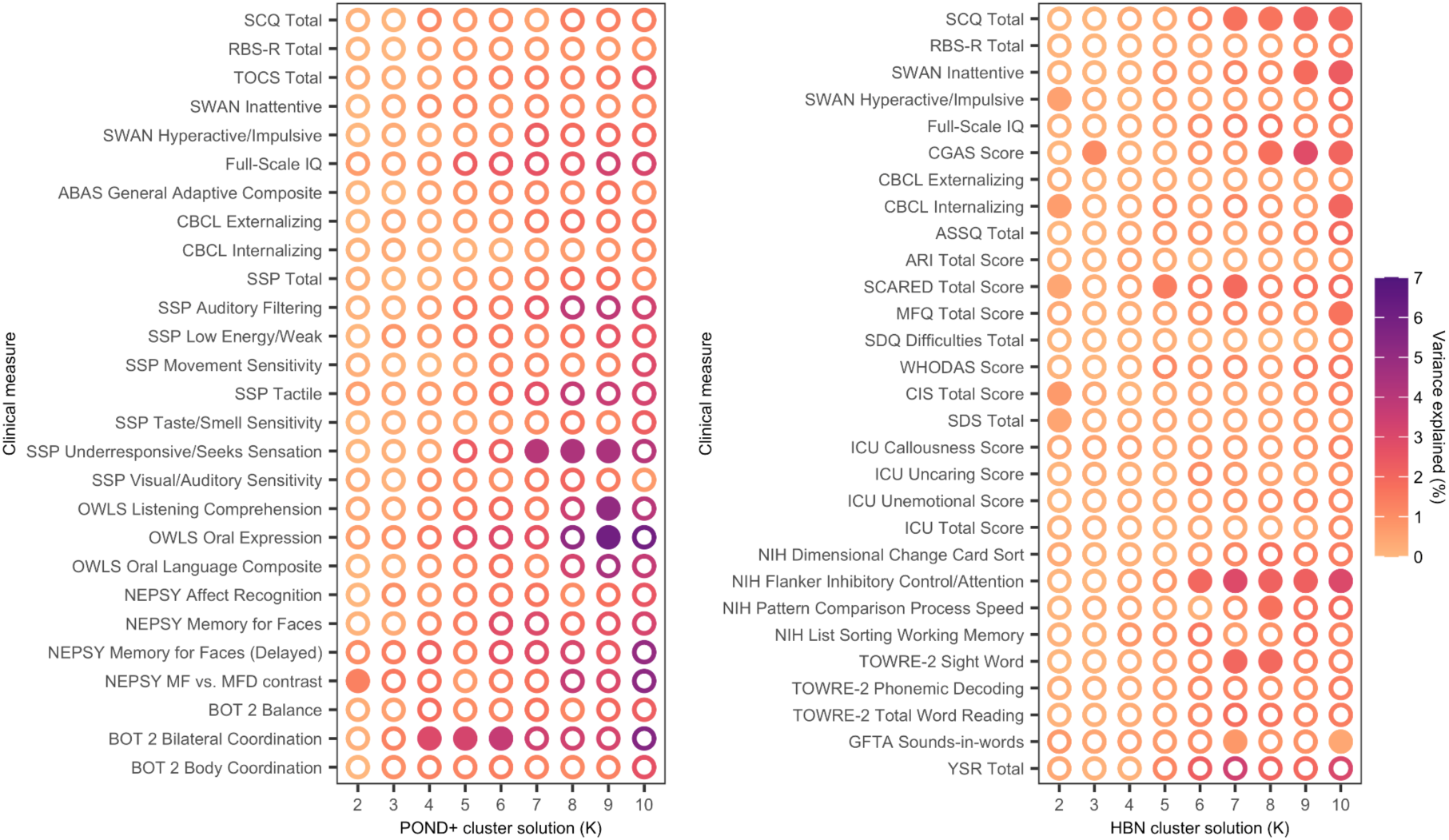
Human clusters are weakly associated with clinical and cognitive scores. Analyses of cognitive and clinical measures for POND+ (left) and HBN (right) reveal that some scores vary significantly between clusters in certain solutions (filled circles). However, these significant results do not survive correction for multiple comparisons. The associations between clusters and scores were evaluated using Kruskal-Wallis tests for each measure and each K. The resulting p-values were corrected for multiple comparisons across measures for each K using the Benjamini-Hochberg procedure^20^.

## Supplementary Tables

**Supplementary Table 1.**
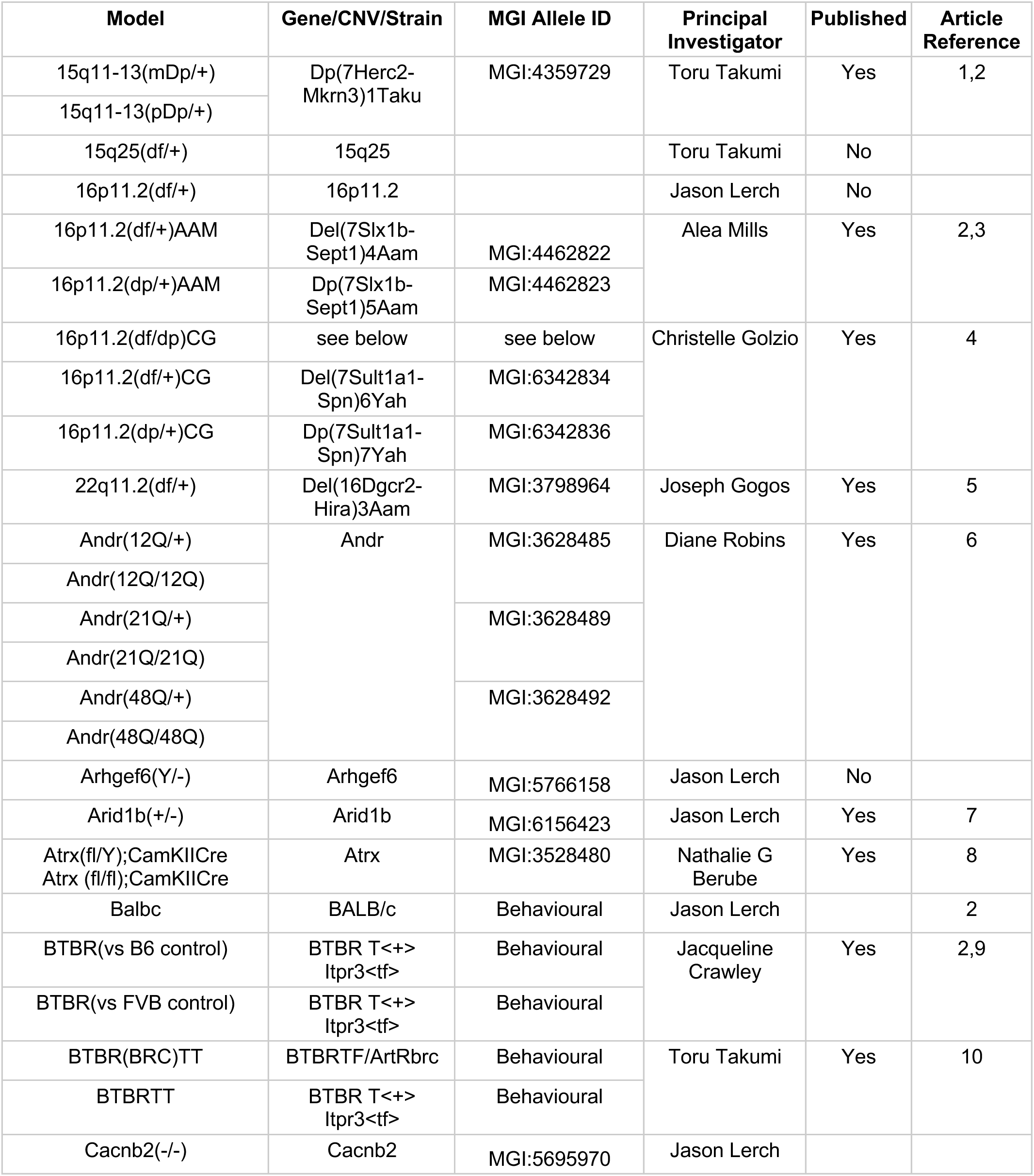
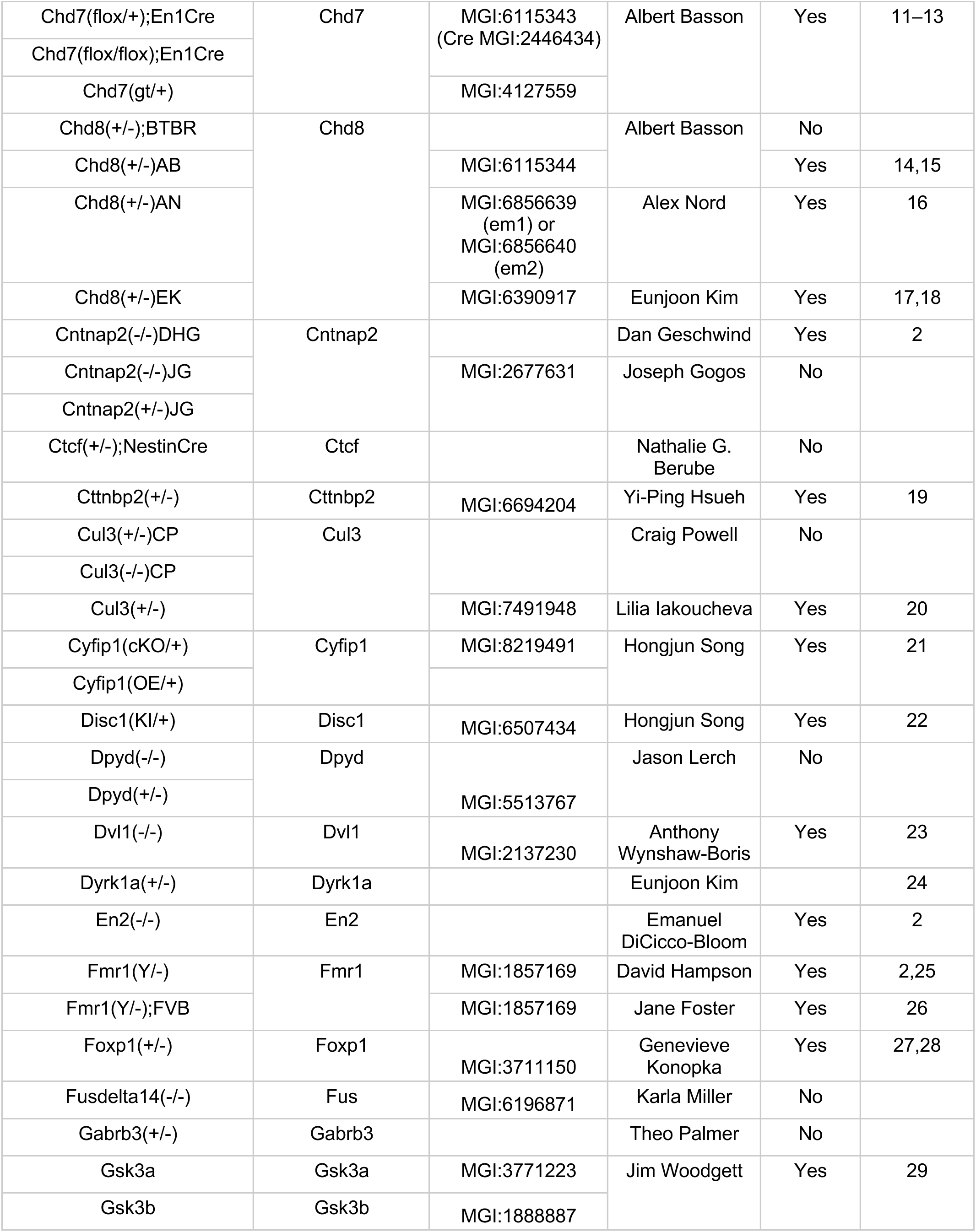
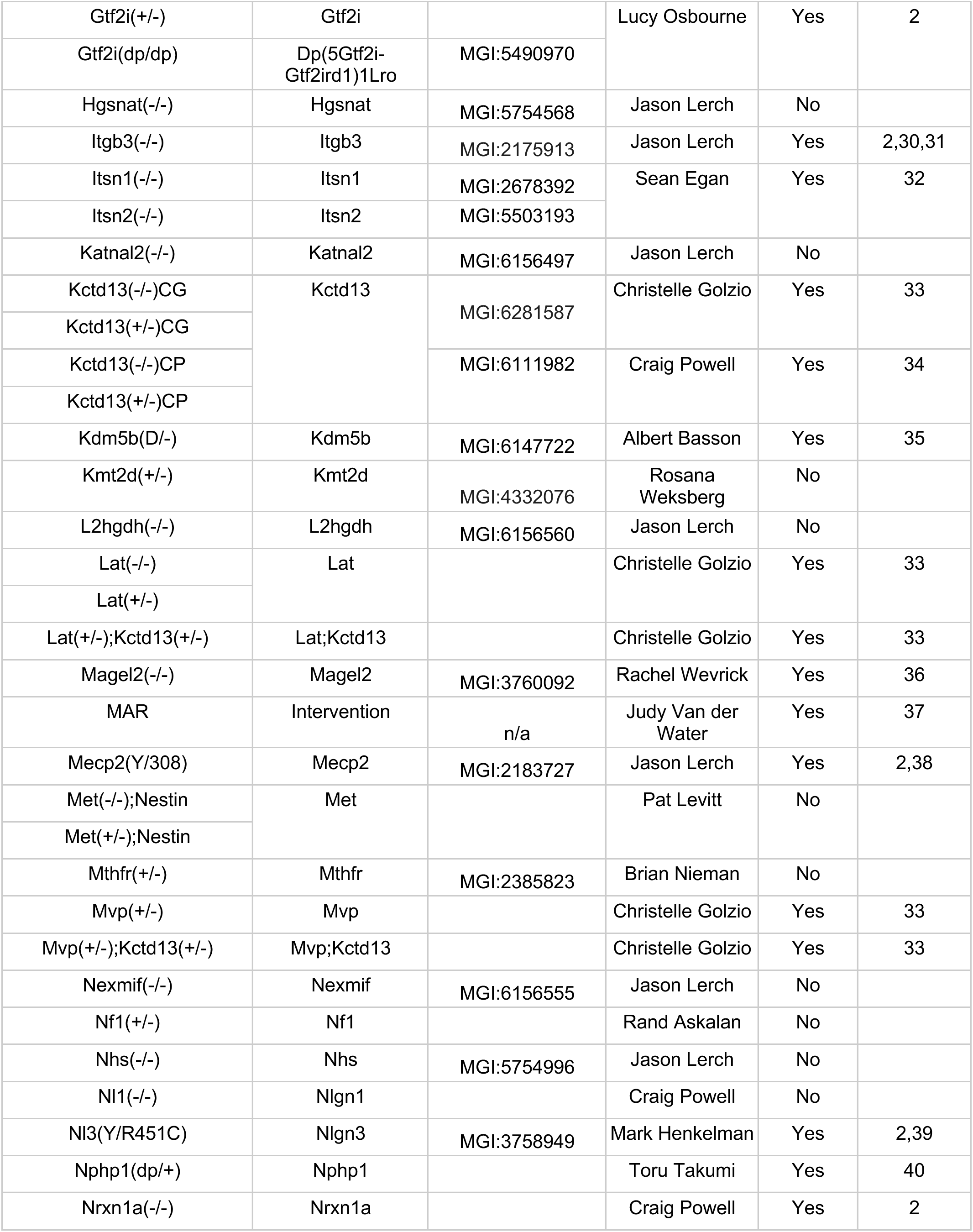
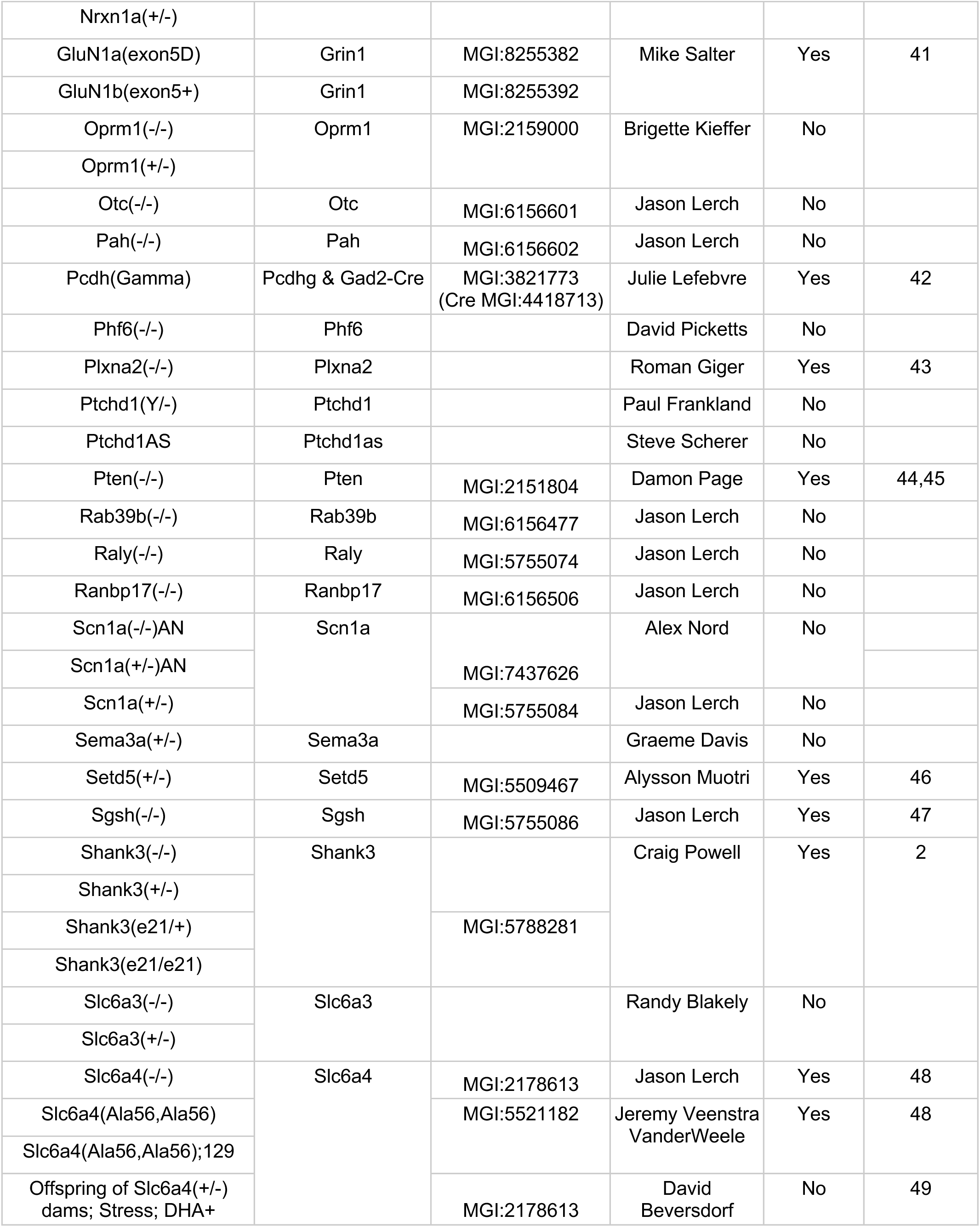
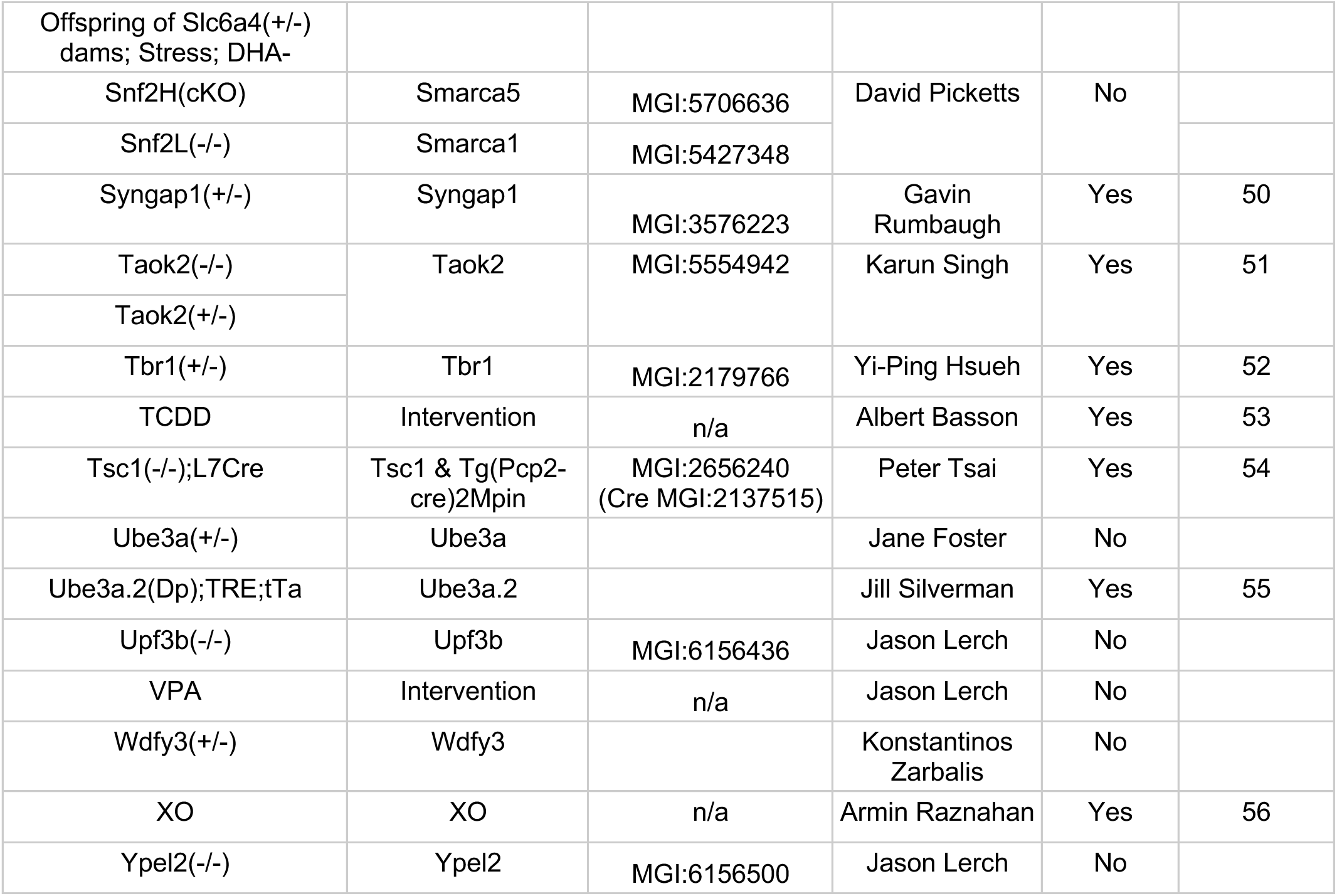
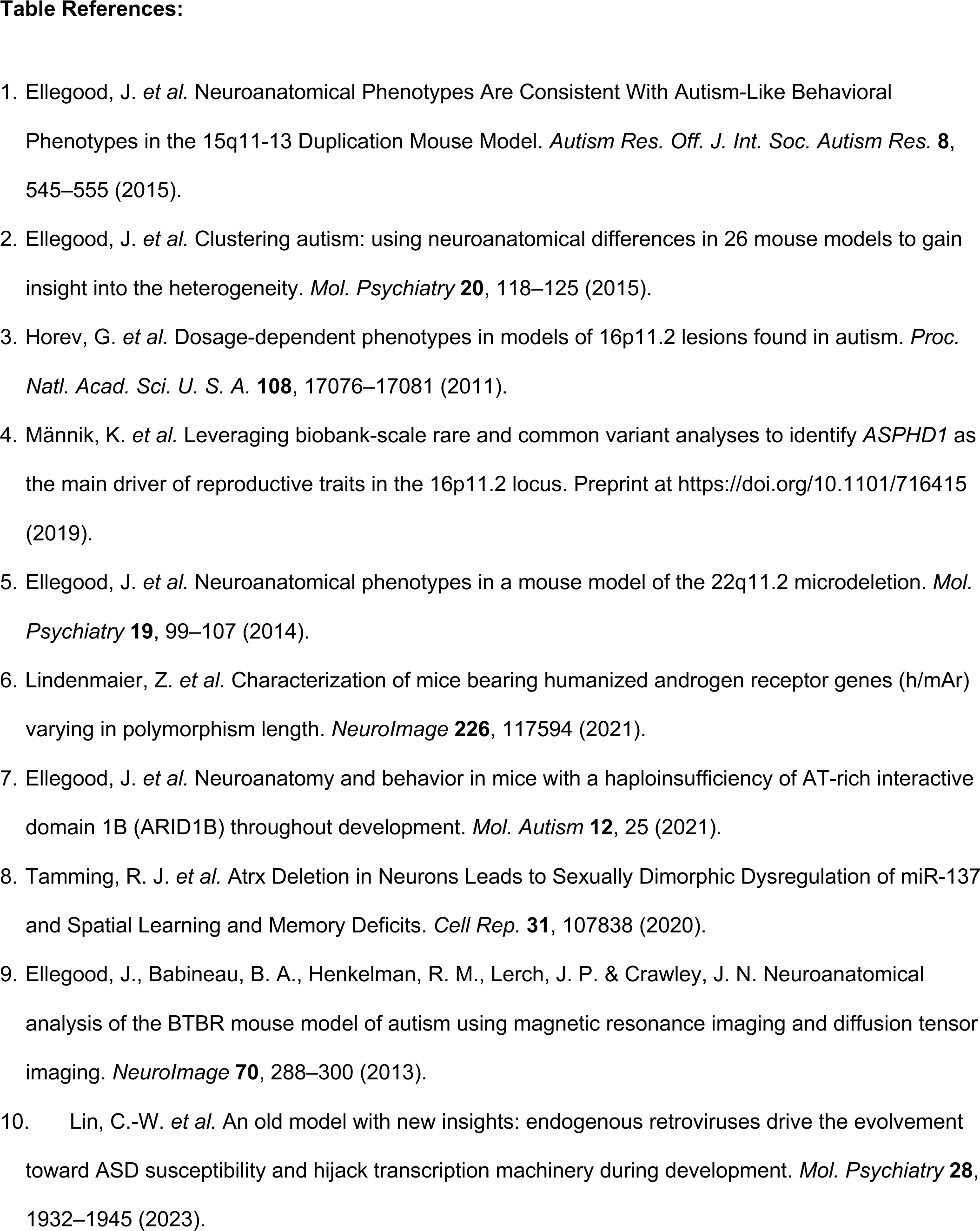
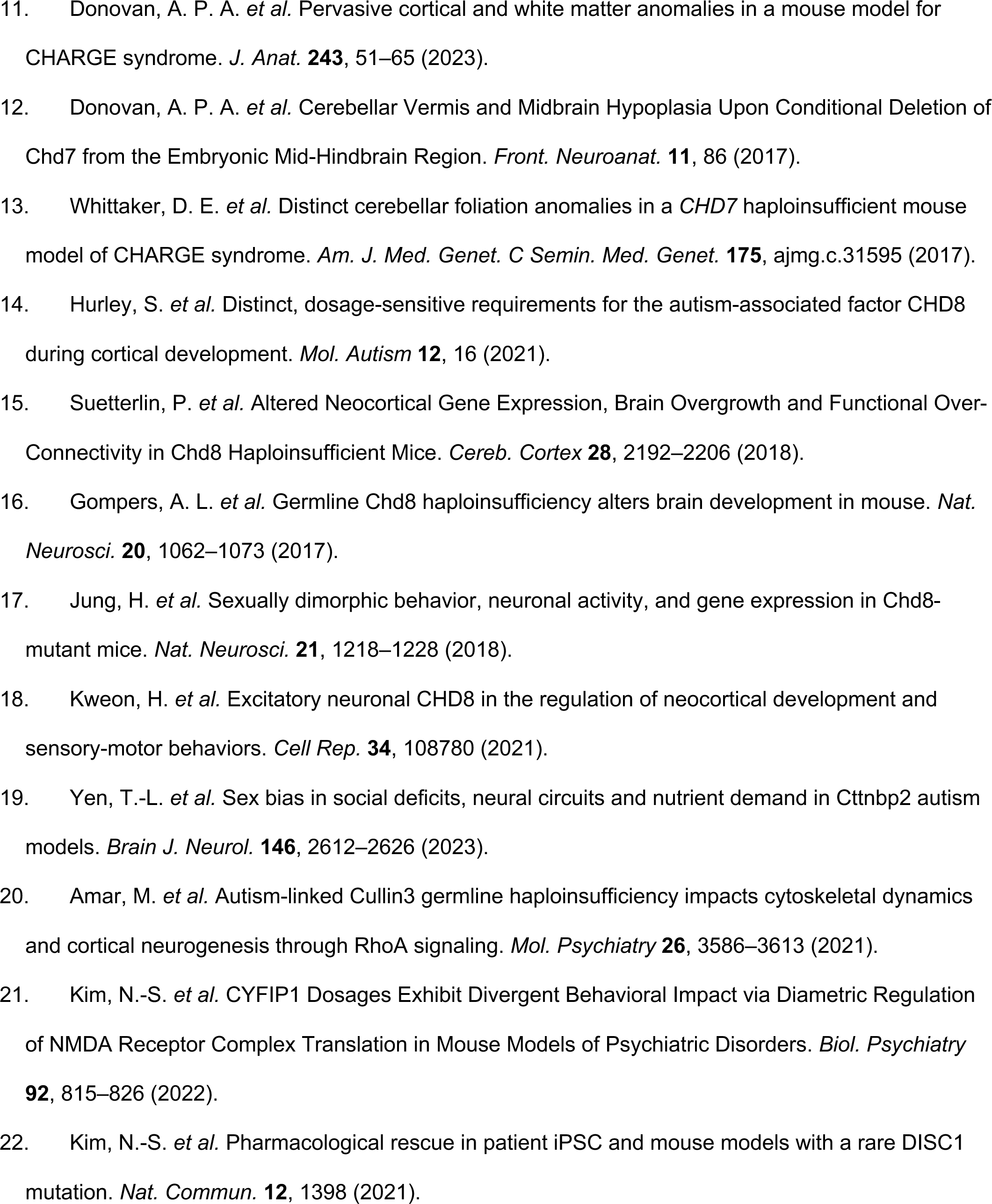
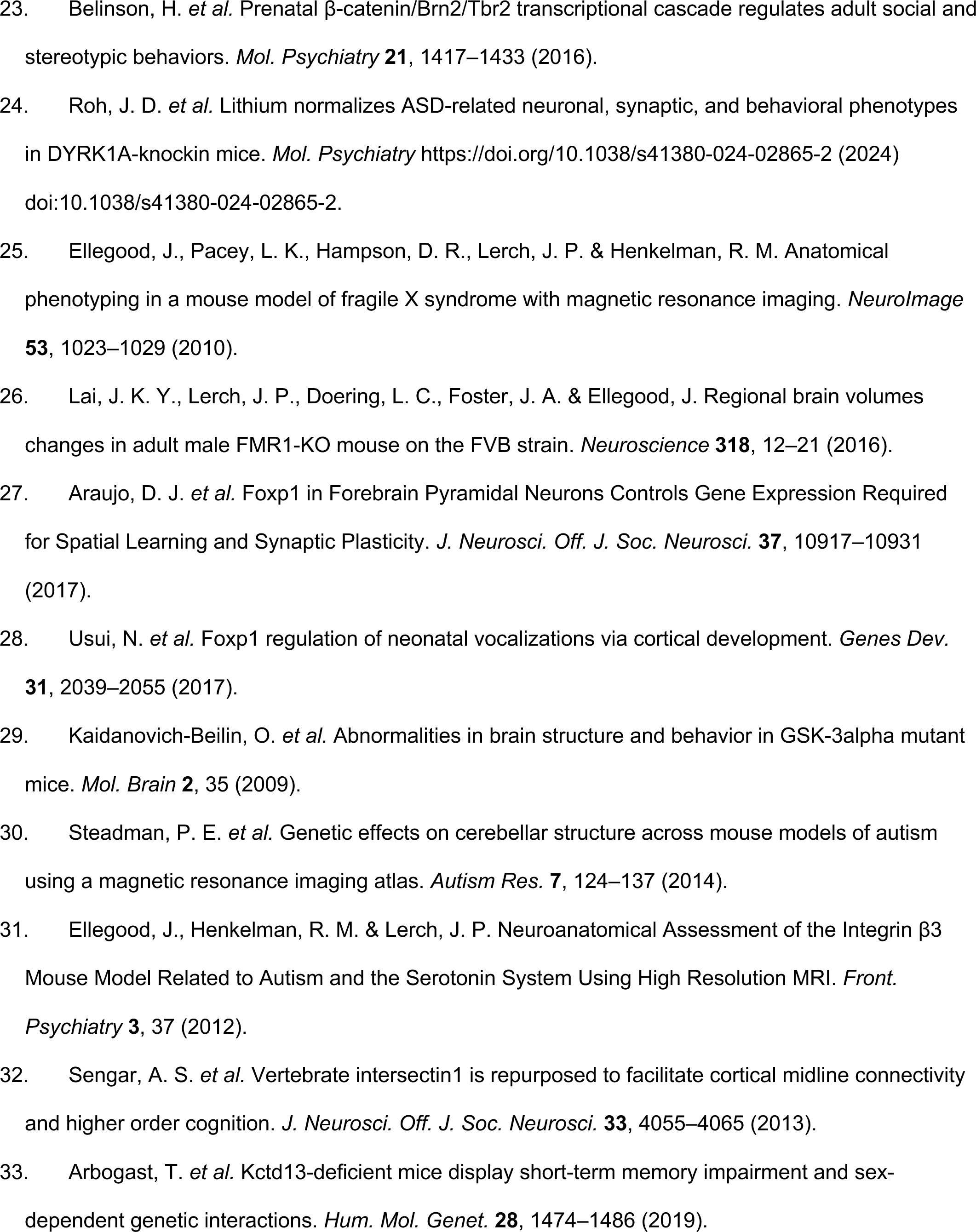
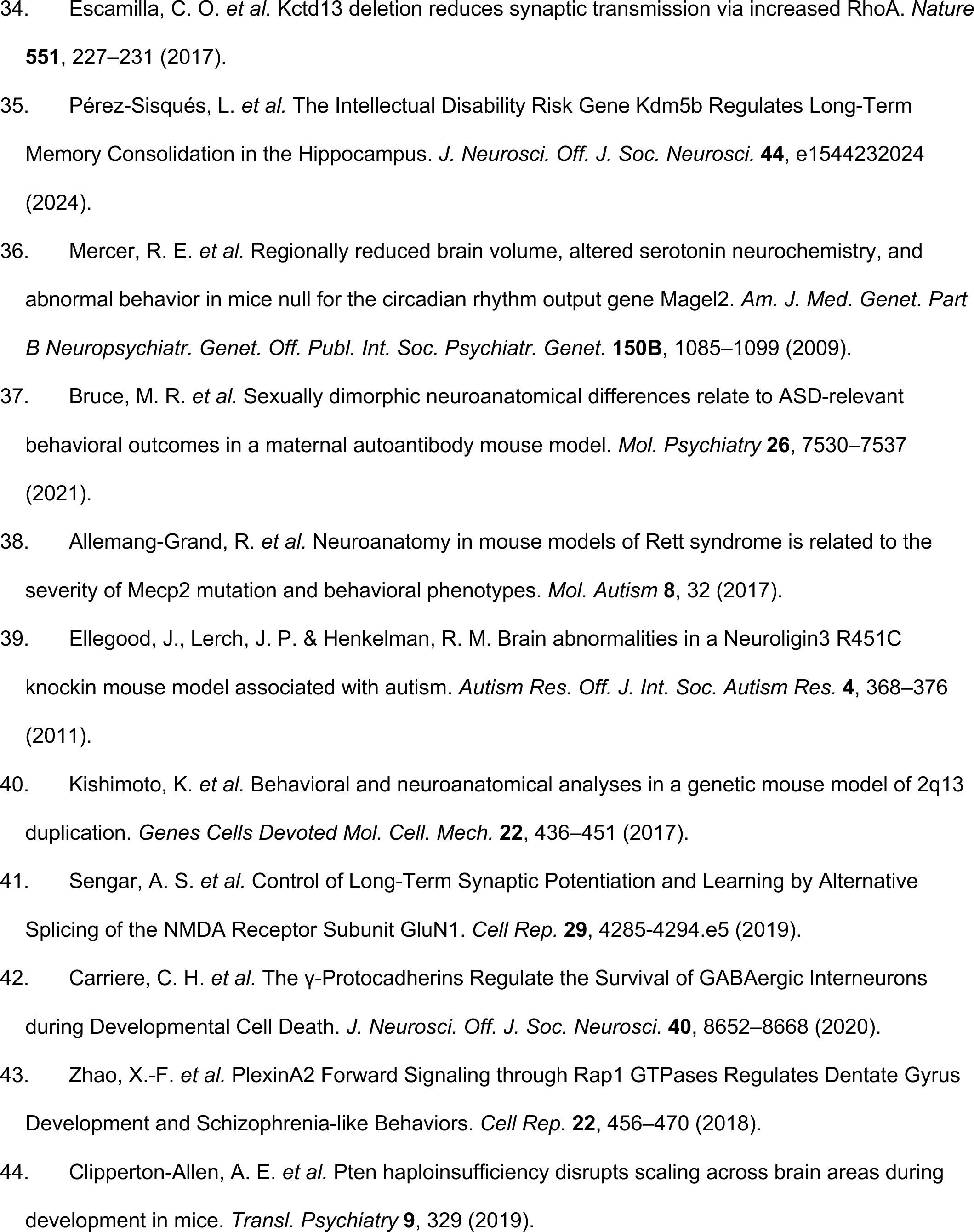
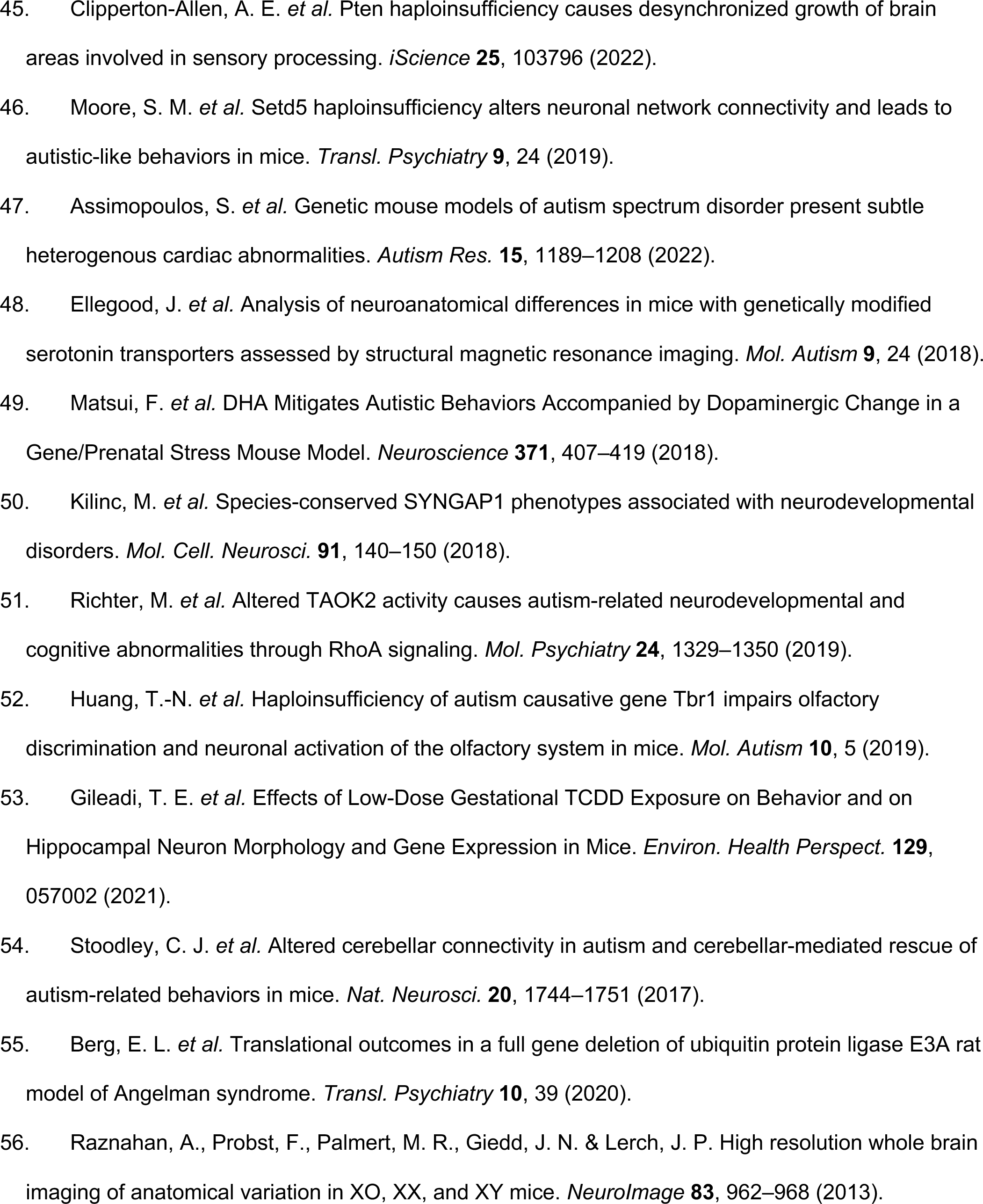
Comprehensive list of mouse models.

**Supplementary Table 2.**
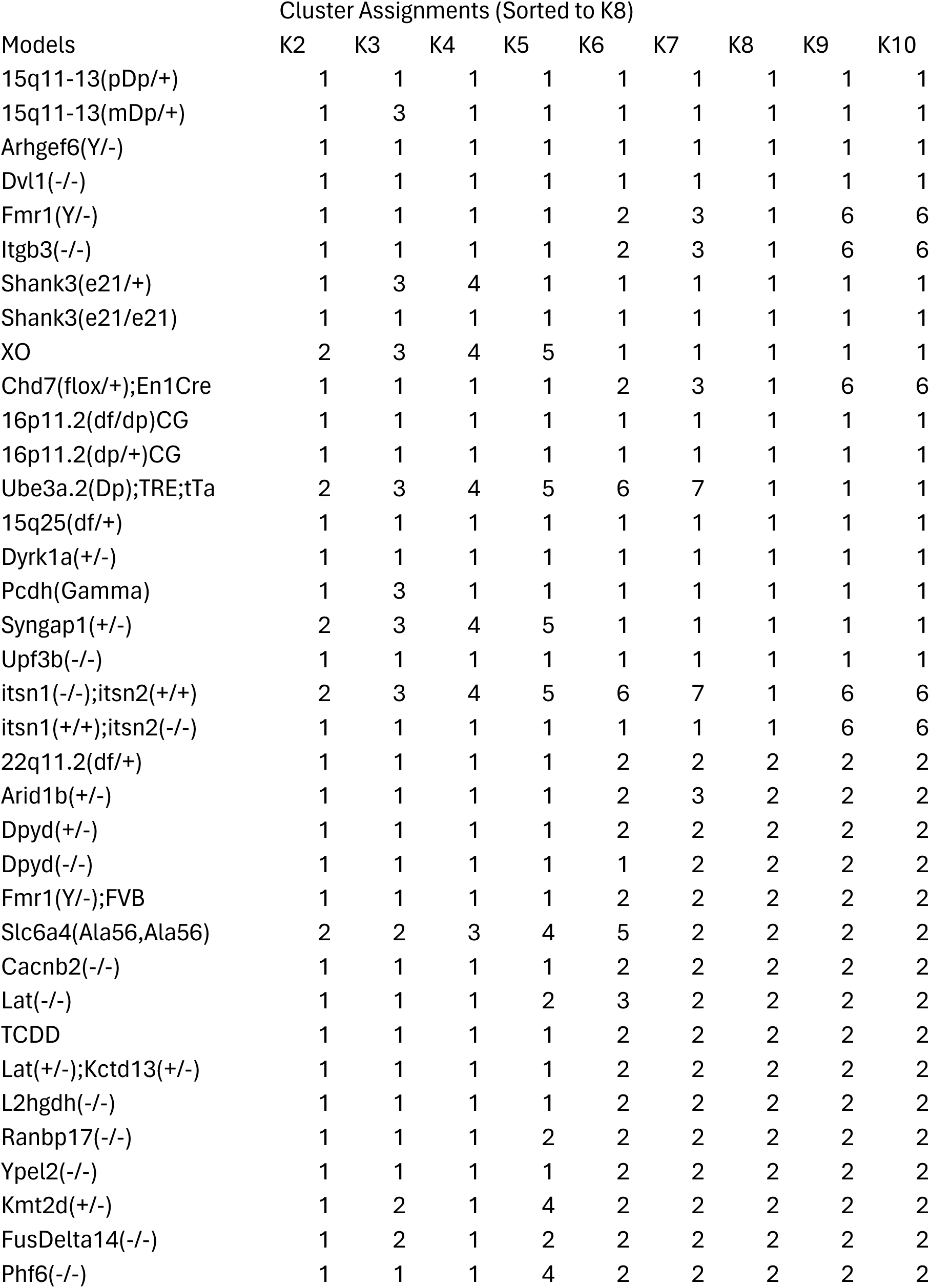
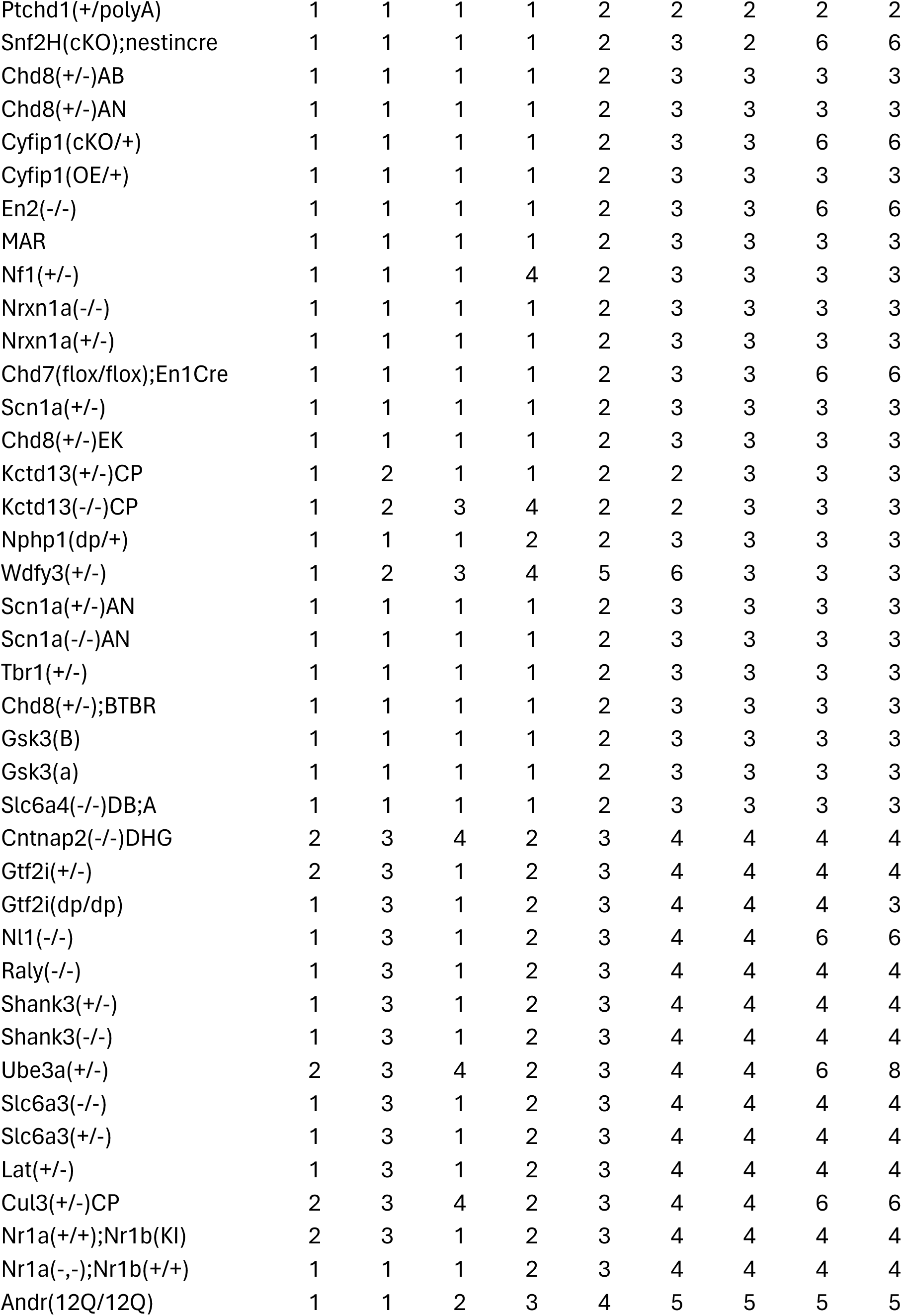
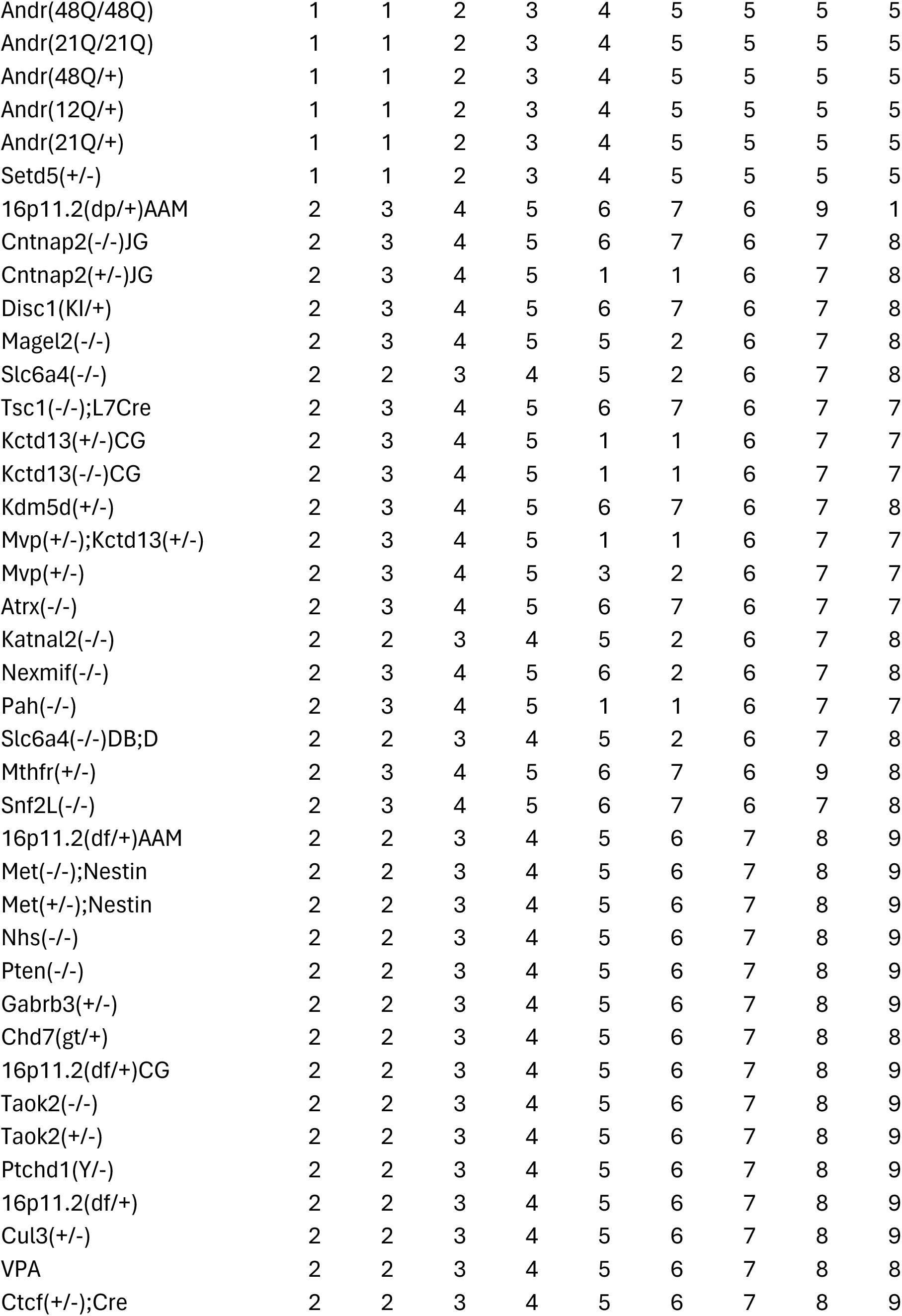
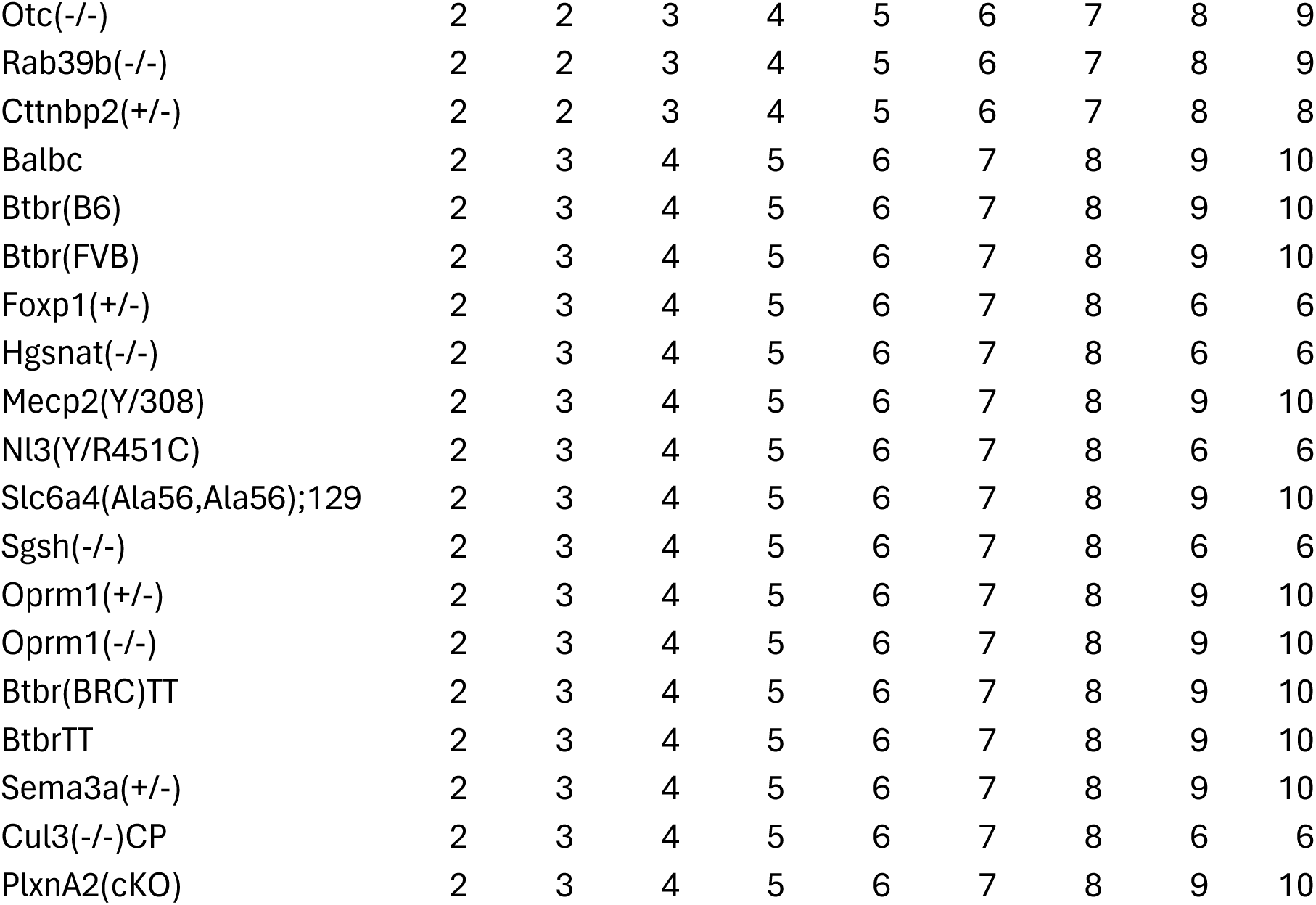
Listing of Mouse Models and Cluster Assignment from K=2-10 Supplementary.

**Supplementary Table 3.** Reactome Pathway List.

| **<u>Pathway Name</u>** | **<u>Reactome Release Information</u>** | **<u>Acronym</u>** |
| --- | --- | --- |
| Signaling by Nuclear Receptors | SIGNALING BY NUCLEAR<br>RECEPTORS%REACTOME%R-HSA-9006931.7 | SNR |
| Cellular responses to stress | CELLULAR RESPONSES TO<br>STRESS%REACTOME%R-HSA-2262752.13 | CRS |
| Adaptive Immune System | ADAPTIVE IMMUNE SYSTEM%REACTOME%R-HSA-1280218.7 | AIS |
| Signaling by Receptor Tyrosine<br>Kinases | SIGNALING BY RECEPTOR TYROSINE<br>KINASES%REACTOME%R-HSA-9006934.8 | SRTK |
| MAPK family signaling<br>cascades | MAPK FAMILY SIGNALING CASCADES%REACTOME<br>DATABASE ID RELEASE 92%5683057 | MAPK |
| Intracellular signaling by<br>second messengers | INTRACELLULAR SIGNALING BY SECOND<br>MESSENGERS%REACTOME DATABASE ID RELEASE<br>92%9006925 | SSM |
| Cytokine Signaling in Immune<br>System | CYTOKINE SIGNALING IN IMMUNE<br>SYSTEM%REACTOME DATABASE ID RELEASE<br>92%1280215 | CSIS |
| Protein-protein interactions at<br>synapses | PROTEIN-PROTEIN INTERACTIONS AT<br>SYNAPSES%REACTOME%R-HSA-6794362.6 | PPIS |
| Transmission across Chemical Synapses | TRANSMISSION ACROSS CHEMICAL SYNAPSES%REACTOME DATABASE ID RELEASE 92%112315 | TACS |
| RNA Polymerase II Transcription | RNA POLYMERASE II TRANSCRIPTION%REACTOME%R-HSA-73857.7 | RPII |
| Epigenetic regulation of gene expression | EPIGENETIC REGULATION OF GENE EXPRESSION%REACTOME DATABASE ID RELEASE 92%212165 | ERGE |
| Chromatin modifying enzymes | CHROMATIN MODIFYING ENZYMES%REACTOME%R-HSA-3247509.6 | CME |
| Signaling by WNT | SIGNALING BY WNT%REACTOME DATABASE ID RELEASE 92%195721 | WNT |
| Nervous system development | NERVOUS SYSTEM DEVELOPMENT%REACTOME DATABASE ID RELEASE 92%9675108 | NSD |
| Signaling by Rho GTPases, Miro GTPases and RHOBTB3 | SIGNALING BY RHO GTPASES, MIRO GTPASES AND RHOBTB3%REACTOME%R-HSA-9716542.4 | GTPases |

## References and Notes

1. A. Kushki, E. Anagnostou, C. Hammill, P. Duez, J. Brian, A. Iaboni, R. Schachar, J. Crosbie, P. Arnold, J. P. Lerch, Examining overlap and homogeneity in ASD, ADHD, and OCD: a data-driven, diagnosis-agnostic approach. Transl. Psychiatry 9, 318 (2019).

2. M. M. Vandewouw, J. Brian, J. Crosbie, R. J. Schachar, A. Iaboni, S. Georgiades, R. Nicolson, E. Kelley, M. Ayub, J. Jones, Identifying replicable subgroups in neurodevelopmental conditions using resting-state functional magnetic resonance imaging data. *JAMA Netw*. Open 6, e232066–e232066 (2023).

3. M. D. Hettwer, S. Larivière, B. Y. Park, O. A. van den Heuvel, L. Schmaal, O. A. Andreassen, C. R. K. Ching, M. Hoogman, J. Buitelaar, D. van Rooij, D. J. Veltman, D. J. Stein, B. Franke, T. G. M. van Erp, ENIGMA ADHD Working Group, ENIGMA Autism Working Group, ENIGMA Bipolar Disorder Working Group, ENIGMA Major Depression Working Group, ENIGMA OCD Working Group, ENIGMA Schizophrenia Working Group, N. Jahanshad, P. M. Thompson, S. I. Thomopoulos, R. a. I. Bethlehem, B. C. Bernhardt, S. B. Eickhoff, S. L. Valk, Coordinated cortical thickness alterations across six neurodevelopmental and psychiatric disorders. Nat. Commun. 13, 6851 (2022).

4. M. Zarrei, C. L. Burton, W. Engchuan, E. J. Young, E. J. Higginbotham, J. R. MacDonald, B. Trost, A. J. S. Chan, S. Walker, S. Lamoureux, T. Heung, B. A. Mojarad, B. Kellam, T. Paton, M. Faheem, K. Miron, C. Lu, T. Wang, K. Samler, X. Wang, G. Costain, N. Hoang, G. Pellecchia, J. Wei, R. V. Patel, B. Thiruvahindrapuram, M. Roifman, D. Merico, T. Goodale, I. Drmic, M. Speevak, J. L. Howe, R. K. C. Yuen, J. A. Buchanan, J. A. S. Vorstman, C. R. Marshall, R. F. Wintle, D. R. Rosenberg, G. L. Hanna, M. Woodbury-Smith, C. Cytrynbaum, L. Zwaigenbaum, M. Elsabbagh, J. Flanagan, B. A. Fernandez, M. T. Carter, P. Szatmari, W. Roberts, J. Lerch, X. Liu, R. Nicolson, S. Georgiades, R. Weksberg, P. D. Arnold, A. S. Bassett, J. Crosbie, R. Schachar, D. J. Stavropoulos, E. Anagnostou, S. W. Scherer, A large data resource of genomic copy number variation across neurodevelopmental disorders. NPJ Genomic Med. 4, 26 (2019).

5. H. Cao, J. Wang, A. Baranova, F. Zhang, Classifying major mental disorders genetically. Prog. Neuropsychopharmacol. Biol. Psychiatry 112, 110410 (2022).

6. J. Cabana-Domínguez, B. Torrico, A. Reif, N. Fernàndez-Castillo, B. Cormand, Comprehensive exploration of the genetic contribution of the dopaminergic and serotonergic pathways to psychiatric disorders. Transl. Psychiatry 12, 11 (2022).

7. The Brainstorm Consortium, V. Anttila, B. Bulik-Sullivan, H. K. Finucane, R. K. Walters, J. Bras, L. Duncan, V. Escott-Price, G. J. Falcone, P. Gormley, R. Malik, N. A. Patsopoulos, S. Ripke, Z. Wei, D. Yu, P. H. Lee, P. Turley, B. Grenier-Boley, V. Chouraki, Y. Kamatani, C. Berr, L. Letenneur, D. Hannequin, P. Amouyel, A. Boland, J.-F. Deleuze, E. Duron, B. N. Vardarajan, C. Reitz, A. M. Goate, M. J. Huentelman, M. I. Kamboh, E. B. Larson, E. Rogaeva, P. St George-Hyslop, H. Hakonarson, W. A. Kukull, L. A. Farrer, L. L. Barnes, T. G. Beach, F. Y. Demirci, E. Head, C. M. Hulette, G. A. Jicha, J. S. K. Kauwe, J. A. Kaye, J. B. Leverenz, A. I. Levey, A. P. Lieberman, V. S. Pankratz, W. W. Poon, J. F. Quinn, A. J. Saykin, L. S. Schneider, A. G. Smith, J. A. Sonnen, R. A. Stern, V. M. Van Deerlin, L. J. Van Eldik, D. Harold, G. Russo, D. C. Rubinsztein, A. Bayer, M. Tsolaki, P. Proitsi, N. C. Fox, H. Hampel, M. J. Owen, S. Mead, P. Passmore, K. Morgan, M. M. Nöthen, J. M. Schott, M. Rossor, M. K. Lupton, P. Hoffmann, J. Kornhuber, B. Lawlor, A. McQuillin, A. Al-Chalabi, J. C. Bis, A. Ruiz, M. Boada, S. Seshadri, A. Beiser, K. Rice, S. J. Van Der Lee, P. L. De Jager, D. H. Geschwind, M. Riemenschneider, S. Riedel-Heller, J. I. Rotter, G. Ransmayr, B. T. Hyman, C. Cruchaga, M. Alegret, B. Winsvold, P. Palta, K.-H. Farh, E. Cuenca-Leon, N. Furlotte, T. Kurth, L. Ligthart, G. M. Terwindt, T. Freilinger, C. Ran, S. D. Gordon, G. Borck, H. H. H. Adams, T. Lehtimäki, J. Wedenoja, J. E. Buring, M. Schürks, M. Hrafnsdottir, J.-J. Hottenga, B. Penninx, V. Artto, M. Kaunisto, S. Vepsäläinen, N. G. Martin, G. W. Montgomery, M. I. Kurki, E. Hämäläinen, H. Huang, J. Huang, C. Sandor, C. Webber, B. Muller-Myhsok, S. Schreiber, V. Salomaa, E. Loehrer, H. Göbel, A. Macaya, P. Pozo-Rosich, T. Hansen, T. Werge, J. Kaprio, A. Metspalu, C. Kubisch, M. D. Ferrari, A. C. Belin, A. M. J. M. Van Den Maagdenberg, J.-A. Zwart, D. Boomsma, N. Eriksson, J. Olesen, D. I. Chasman, D. R. Nyholt, R. Anney, A. Avbersek, L. Baum, S. Berkovic, J. Bradfield, R. J. Buono, C. B. Catarino, P. Cossette, P. De Jonghe, C. Depondt, D. Dlugos, T. N. Ferraro, J. French, H. Hjalgrim, J. Jamnadas-Khoda, R. Kälviäinen, W. S. Kunz, H. Lerche, C. Leu, D. Lindhout, W. Lo, D. Lowenstein, M. McCormack, R. S. Møller, A. Molloy, P.-W. Ng, K. Oliver, M. Privitera, R. Radtke, A.-K. Ruppert, T. Sander, S. Schachter, C. Schankin, I. Scheffer, S. Schoch, S. M. Sisodiya, P. Smith, M. Sperling, P. Striano, R. Surges, G. N. Thomas, F. Visscher, C. D. Whelan, F. Zara, E. L. Heinzen, A. Marson, F. Becker, H. Stroink, F. Zimprich, T. Gasser, R. Gibbs, P. Heutink, M. Martinez, H. R. Morris, M. Sharma, M. Ryten, K. Y. Mok, S. Pulit, S. Bevan, E. Holliday, J. Attia, T. Battey, G. Boncoraglio, V. Thijs, W.-M. Chen, B. Mitchell, P. Rothwell, P. Sharma, C. Sudlow, A. Vicente, H. Markus, C. Kourkoulis, J. Pera, M. Raffeld, S. Silliman, V. Boraska Perica, L. M. Thornton, L. M. Huckins, N. William Rayner, C. M. Lewis, M. Gratacos, F. Rybakowski, A. Keski-Rahkonen, A. Raevuori, J. I. Hudson, T. Reichborn-Kjennerud, P. Monteleone, A. Karwautz, K. Mannik, J. H. Baker, J. K. O’Toole, S. E. Trace, O. S. P. Davis, S. G. Helder, S. Ehrlich, B. Herpertz-Dahlmann, U. N. Danner, A. A. Van Elburg, M. Clementi, M. Forzan, E. Docampo, J. Lissowska, J. Hauser, A. Tortorella, M. Maj, F. Gonidakis, K. Tziouvas, H. Papezova, Z. Yilmaz, G. Wagner, S. Cohen-Woods, S. Herms, A. Julià, R. Rabionet, D. M. Dick, S. Ripatti, O. A. Andreassen, T. Espeseth, A. J. Lundervold, V. M. Steen, D. Pinto, S. W. Scherer, H. Aschauer, A. Schosser, L. Alfredsson, L. Padyukov, K. A. Halmi, J. Mitchell, M. Strober, A. W. Bergen, W. Kaye, J. P. Szatkiewicz, B. Cormand, J. A. Ramos-Quiroga, C. Sánchez-Mora, M. Ribasés, M. Casas, A. Hervas, M. J. Arranz, J. Haavik, T. Zayats, S. Johansson, N. Williams, J. Elia, A. Dempfle, A. Rothenberger, J. Kuntsi, R. D. Oades, T. Banaschewski, B. Franke, J. K. Buitelaar, A. Arias Vasquez, A. E. Doyle, A. Reif, K.-P. Lesch, C. Freitag, O. Rivero, H. Palmason, M. Romanos, K. Langley, M. Rietschel, S. H. Witt, S. Dalsgaard, A. D. Børglum, I. Waldman, B. Wilmot, N. Molly, C. H. D. Bau, J. Crosbie, R. Schachar, S. K. Loo, J. J. McGough, E. H. Grevet, S. E. Medland, E. Robinson, L. A. Weiss, E. Bacchelli, A. Bailey, V. Bal, A. Battaglia, C. Betancur, P. Bolton, R. Cantor, P. Celestino-Soper, G. Dawson, S. De Rubeis, F. Duque, A. Green, S. M. Klauck, M. Leboyer, P. Levitt, E. Maestrini, S. Mane, D. M.- De-Luca, J. Parr, R. Regan, A. Reichenberg, S. Sandin, J. Vorstman, T. Wassink, E. Wijsman, E. Cook, S. Santangelo, R. Delorme, B. Rogé, T. Magalhaes, D. Arking, T. G. Schulze, R. C. Thompson, J. Strohmaier, K. Matthews, I. Melle, D. Morris, D. Blackwood, A. McIntosh, S. E. Bergen, M. Schalling, S. Jamain, A. Maaser, S. B. Fischer, C. S. Reinbold, J. M. Fullerton, M. Grigoroiu-Serbanescu, J. Guzman-Parra, F. Mayoral, P. R. Schofield, S. Cichon, T. W. Mühleisen, F. Degenhardt, J. Schumacher, M. Bauer, P. B. Mitchell, E. S. Gershon, J. Rice, J. B. Potash, P. P. Zandi, N. Craddock, I. N. Ferrier, M. Alda, G. A. Rouleau, G. Turecki, R. Ophoff, C. Pato, A. Anjorin, E. Stahl, M. Leber, P. M. Czerski, H. J. Edenberg, C. Cruceanu, I. R. Jones, D. Posthuma, T. F. M. Andlauer, A. J. Forstner, F. Streit, B. T. Baune, T. Air, G. Sinnamon, N. R. Wray, D. J. MacIntyre, D. Porteous, G. Homuth, M. Rivera, J. Grove, C. M. Middeldorp, I. Hickie, M. Pergadia, D. Mehta, J. H. Smit, R. Jansen, E. De Geus, E. Dunn, Q. S. Li, M. Nauck, R. A. Schoevers, A. T. Beekman, J. A. Knowles, A. Viktorin, P. Arnold, C. L. Barr, G. Bedoya-Berrio, O. J. Bienvenu, H. Brentani, C. Burton, B. Camarena, C. Cappi, D. Cath, M. Cavallini, D. Cusi, S. Darrow, D. Denys, E. M. Derks, A. Dietrich, T. Fernandez, M. Figee, N. Freimer, G. Gerber, M. Grados, E. Greenberg, G. L. Hanna, A. Hartmann, M. E. Hirschtritt, P. J. Hoekstra, A. Huang, C. Huyser, C. Illmann, M. Jenike, S. Kuperman, B. Leventhal, C. Lochner, G. J. Lyon, F. Macciardi, M. Madruga-Garrido, I. A. Malaty, A. Maras, L. McGrath, E. C. Miguel, P. Mir, G. Nestadt, H. Nicolini, M. S. Okun, A. Pakstis, P. Paschou, J. Piacentini, C. Pittenger, K. Plessen, V. Ramensky, E. M. Ramos, V. Reus, M. A. Richter, M. A. Riddle, M. M. Robertson, V. Roessner, M. Rosário, J. F. Samuels, P. Sandor, D. J. Stein, F. Tsetsos, F. Van Nieuwerburgh, S. Weatherall, J. R. Wendland, T. Wolanczyk, Y. Worbe, G. Zai, F. S. Goes, N. McLaughlin, P. S. Nestadt, H.-J. Grabe, C. Depienne, A. Konkashbaev, N. Lanzagorta, A. Valencia-Duarte, E. Bramon, N. Buccola, W. Cahn, M. Cairns, S. A. Chong, D. Cohen, B. Crespo-Facorro, J. Crowley, M. Davidson, L. DeLisi, T. Dinan, G. Donohoe, E. Drapeau, J. Duan, L. Haan, D. Hougaard, S. Karachanak-Yankova, A. Khrunin, J. Klovins, V. Kučinskas, J. Lee Chee Keong, S. Limborska, C. Loughland, J. Lönnqvist, B. Maher, M. Mattheisen, C. McDonald, K. C. Murphy, R. Murray, I. Nenadic, J. Van Os, C. Pantelis, M. Pato, T. Petryshen, D. Quested, P. Roussos, A. R. Sanders, U. Schall, S. G. Schwab, K. Sim, H.-C. So, E. Stögmann, M. Subramaniam, D. Toncheva, J. Waddington, J. Walters, M. Weiser, W. Cheng, R. Cloninger, D. Curtis, P. V. Gejman, F. Henskens, M. Mattingsdal, S.-Y. Oh, R. Scott, B. Webb, G. Breen, C. Churchhouse, C. M. Bulik, M. Daly, M. Dichgans, S. V. Faraone, R. Guerreiro, P. Holmans, K. S. Kendler, B. Koeleman, C. A. Mathews, A. Price, J. Scharf, P. Sklar, J. Williams, N. W. Wood, C. Cotsapas, A. Palotie, J. W. Smoller, P. Sullivan, J. Rosand, A. Corvin, B. M. Neale, Analysis of shared heritability in common disorders of the brain. Science 360, eaap8757 (2018).

8. L. M. Alexander, J. Escalera, L. Ai, C. Andreotti, K. Febre, A. Mangone, N. Vega-Potler, N. Langer, A. Alexander, M. Kovacs, S. Litke, B. O’Hagan, J. Andersen, B. Bronstein, A. Bui, M. Bushey, H. Butler, V. Castagna, N. Camacho, E. Chan, D. Citera, J. Clucas, S. Cohen, S. Dufek, M. Eaves, B. Fradera, J. Gardner, N. Grant-Villegas, G. Green, C. Gregory, E. Hart, S. Harris, M. Horton, D. Kahn, K. Kabotyanski, B. Karmel, S. P. Kelly, K. Kleinman, B. Koo, E. Kramer, E. Lennon, C. Lord, G. Mantello, A. Margolis, K. R. Merikangas, J. Milham, G. Minniti, R. Neuhaus, A. Levine, Y. Osman, L. C. Parra, K. R. Pugh, A. Racanello, A. Restrepo, T. Saltzman, B. Septimus, R. Tobe, R. Waltz, A. Williams, A. Yeo, F. X. Castellanos, A. Klein, T. Paus, B. L. Leventhal, R. C. Craddock, H. S. Koplewicz, M. P. Milham, An open resource for transdiagnostic research in pediatric mental health and learning disorders. Sci. Data 4, 170181 (2017).

9. D. A. Baribeau, A. Dupuis, T. A. Paton, C. Hammill, S. W. Scherer, R. J. Schachar, P. D. Arnold, P. Szatmari, R. Nicolson, S. Georgiades, J. Crosbie, J. Brian, A. Iaboni, A. Kushki, J. P. Lerch, E. Anagnostou, Structural neuroimaging correlates of social deficits are similar in autism spectrum disorder and attention-deficit/hyperactivity disorder: analysis from the POND Network. Transl. Psychiatry 9, 72 (2019).

10. S. Folstein, M. Rutter, Infantile autism: a genetic study of 21 twin pairs. J. Child Psychol. Psychiatry 18, 297–321 (1977).

11. S. Folstein, M. Rutter, Genetic influences and infantile autism. Nature 265, 726–728 (1977).

12. J. M. Fu, F. K. Satterstrom, M. Peng, H. Brand, R. L. Collins, S. Dong, B. Wamsley, L. Klei, L. Wang, S. P. Hao, C. R. Stevens, C. Cusick, M. Babadi, E. Banks, B. Collins, S. Dodge, S. B. Gabriel, L. Gauthier, S. K. Lee, L. Liang, A. Ljungdahl, B. Mahjani, L. Sloofman, A. N. Smirnov, M. Barbosa, C. Betancur, A. Brusco, B. H. Y. Chung, E. H. Cook, M. L. Cuccaro, E. Domenici, G. B. Ferrero, J. J. Gargus, G. E. Herman, I. Hertz-Picciotto, P. Maciel, D. S. Manoach, M. R. Passos-Bueno, A. M. Persico, A. Renieri, J. S. Sutcliffe, F. Tassone, E. Trabetti, G. Campos, S. Cardaropoli, D. Carli, M. C. Y. Chan, C. Fallerini, E. Giorgio, A. C. Girardi, E. Hansen-Kiss, S. L. Lee, C. Lintas, Y. Ludena, R. Nguyen, L. Pavinato, M. Pericak-Vance, I. N. Pessah, R. J. Schmidt, M. Smith, C. I. S. Costa, S. Trajkova, J. Y. T. Wang, M. H. C. Yu, Autism Sequencing Consortium (ASC), Broad Institute Center for Common Disease Genomics (Broad-CCDG), iPSYCH-BROAD Consortium, D. J. Cutler, S. De Rubeis, J. D. Buxbaum, M. J. Daly, B. Devlin, K. Roeder, S. J. Sanders, M. E. Talkowski, Rare coding variation provides insight into the genetic architecture and phenotypic context of autism. Nat. Genet. 54, 1320–1331 (2022).

13. F. K. Satterstrom, J. A. Kosmicki, J. Wang, M. S. Breen, S. De Rubeis, J.-Y. An, M. Peng, R. Collins, J. Grove, L. Klei, C. Stevens, J. Reichert, M. S. Mulhern, M. Artomov, S. Gerges, B. Sheppard, X. Xu, A. Bhaduri, U. Norman, H. Brand, G. Schwartz, R. Nguyen, E. E. Guerrero, C. Dias, Autism Sequencing Consortium, iPSYCH-Broad Consortium, C. Betancur, E. H. Cook, L. Gallagher, M. Gill, J. S. Sutcliffe, A. Thurm, M. E. Zwick, A. D. Børglum, M. W. State, A. E. Cicek, M. E. Talkowski, D. J. Cutler, B. Devlin, S. J. Sanders, K. Roeder, M. J. Daly, J. D. Buxbaum, Large-Scale Exome Sequencing Study Implicates Both Developmental and Functional Changes in the Neurobiology of Autism. Cell 180, 568–584.e23 (2020).

14. B. Trost, B. Thiruvahindrapuram, A. J. S. Chan, W. Engchuan, E. J. Higginbotham, J. L. Howe, L. O. Loureiro, M. S. Reuter, D. Roshandel, J. Whitney, M. Zarrei, M. Bookman, C. Somerville, R. Shaath, M. Abdi, E. Aliyev, R. V. Patel, T. Nalpathamkalam, G. Pellecchia, O. Hamdan, G. Kaur, Z. Wang, J. R. MacDonald, J. Wei, W. W. L. Sung, S. Lamoureux, N. Hoang, T. Selvanayagam, N. Deflaux, M. Geng, S. Ghaffari, J. Bates, E. J. Young, Q. Ding, C. Shum, L. D’Abate, C. A. Bradley, A. Rutherford, V. Aguda, B. Apresto, N. Chen, S. Desai, X. Du, M. L. Y. Fong, S. Pullenayegum, K. Samler, T. Wang, K. Ho, T. Paton, S. L. Pereira, J.-A. Herbrick, R. F. Wintle, J. Fuerth, J. Noppornpitak, H. Ward, P. Magee, A. Al Baz, U. Kajendirarajah, S. Kapadia, J. Vlasblom, M. Valluri, J. Green, V. Seifer, M. Quirbach, O. Rennie, E. Kelley, N. Masjedi, C. Lord, M. J. Szego, M. H. Zawati, M. Lang, L. J. Strug, C. R. Marshall, G. Costain, K. Calli, A. Iaboni, A. Yusuf, P. Ambrozewicz, L. Gallagher, D. G. Amaral, J. Brian, M. Elsabbagh, S. Georgiades, D. S. Messinger, S. Ozonoff, J. Sebat, C. Sjaarda, I. M. Smith, P. Szatmari, L. Zwaigenbaum, A. Kushki, T. W. Frazier, J. A. S. Vorstman, K. A. Fakhro, B. A. Fernandez, M. E. S. Lewis, R. Weksberg, M. Fiume, R. K. C. Yuen, E. Anagnostou, N. Sondheimer, D. Glazer, D. M. Hartley, S. W. Scherer, Genomic architecture of autism from comprehensive whole-genome sequence annotation. Cell 185, 4409–4427.e18 (2022).

15. D. H. Geschwind, J. Flint, Genetics and genomics of psychiatric disease. Science 349, 1489–1494 (2015).

16. B. S. Abrahams, D. H. Geschwind, Connecting genes to brain in the autism spectrum disorders. Arch. Neurol. 67, 395–399 (2010).

17. I. Iossifov, B. J. O’roak, S. J. Sanders, M. Ronemus, N. Krumm, D. Levy, H. A. Stessman, K. T. Witherspoon, L. Vives, K. E. Patterson, The contribution of de novo coding mutations to autism spectrum disorder. Nature 515, 216–221 (2014).

18. D. H. Geschwind, Gene hunting in autism spectrum disorder: on the path to precision medicine. Lancet Neurol. 14, 1109–1120 (2015).

19. R. Grzadzinski, M. Huerta, C. Lord, DSM-5 and autism spectrum disorders (ASDs): an opportunity for identifying ASD subtypes. Mol. Autism 4, 1–6 (2013).

20. H. Cholemkery, J. Medda, T. Lempp, C. M. Freitag, Classifying autism spectrum disorders by ADI-R: subtypes or severity gradient? J. Autism Dev. Disord. 46, 2327–2339 (2016).

21. A. Kushki, R. E. Cardy, S. Panahandeh, M. Malihi, C. Hammill, J. Brian, A. Iaboni, M. J. Taylor, R. Schachar, J. Crosbie, Cross-diagnosis structural correlates of autistic-like social communication differences. Cereb. Cortex 31, 5067–5076 (2021).

22. G. R. Jacobs, A. N. Voineskos, C. Hawco, L. Stefanik, N. J. Forde, E. W. Dickie, M.-C. Lai, P. Szatmari, R. Schachar, J. Crosbie, Integration of brain and behavior measures for identification of data-driven groups cutting across children with ASD, ADHD, or OCD. Neuropsychopharmacology 46, 643–653 (2021).

23. A. M. Buch, P. E. Vértes, J. Seidlitz, S. H. Kim, L. Grosenick, C. Liston, Molecular and network-level mechanisms explaining individual differences in autism spectrum disorder. Nat. Neurosci. 26, 650–663 (2023).

24. J. Ellegood, E. Anagnostou, B. A. Babineau, J. N. Crawley, L. Lin, M. Genestine, E. DiCicco-Bloom, J. K. Y. Lai, J. A. Foster, O. Peñagarikano, Clustering autism: using neuroanatomical differences in 26 mouse models to gain insight into the heterogeneity. Mol. Psychiatry 20, 118–125 (2015).

25. M. Pagani, V. Zerbi, S. Gini, F. G. Alvino, A. Banerjee, A. Barberis, M. A. Basson, Y. Bozzi, A. Galbusera, J. Ellegood, M. Fagiolini, J. P. Lerch, M. Matteoli, C. Montani, D. Pozzi, G. Provenzano, M. L. Scattoni, N. Wenderoth, T. Xu, M. V. Lombardo, M. P. Milham, A. Di Martino, A. Gozzi, Autism subtypes identified using cross-species functional connectivity analyses. Nat. Neurosci. 29, 1476–1487 (2026).

26. D. Sussman, R. C. Leung, V. M. Vogan, W. Lee, S. Trelle, S. Lin, D. B. Cassel, M. M. Chakravarty, J. P. Lerch, E. Anagnostou, M. J. Taylor, The autism puzzle: Diffuse but not pervasive neuroanatomical abnormalities in children with ASD. NeuroImage Clin. 8, 170–179 (2015).

27. A. Beauchamp, Y. Yee, B. C. Darwin, A. Raznahan, R. B. Mars, J. P. Lerch, Whole-brain comparison of rodent and human brains using spatial transcriptomics. elife 11, e79418 (2022).

28. C. von Mering, M. Huynen, D. Jaeggi, S. Schmidt, P. Bork, B. Snel, STRING: a database of predicted functional associations between proteins. Nucleic Acids Res. 31, 258–261 (2003).

29. D. Szklarczyk, A. L. Gable, D. Lyon, A. Junge, S. Wyder, J. Huerta-Cepas, M. Simonovic, N. T. Doncheva, J. H. Morris, P. Bork, STRING v11: protein–protein association networks with increased coverage, supporting functional discovery in genome-wide experimental datasets. Nucleic Acids Res. 47, D607–D613 (2019).

30. D. Merico, R. Isserlin, O. Stueker, A. Emili, G. D. Bader, Enrichment map: a network-based method for gene-set enrichment visualization and interpretation. PloS One 5, e13984 (2010).

31. E. Ragueneau, C. Gong, P. Sinquin, C. Sevilla, D. Beavers, A. Grentner, J. Griss, G. F. J. Hogue, N. T. Li, L. Matthews, B. May, M. Milacic, H. Mohammadi, R. Petryszak, K. Rothfels, V. Shamovsky, R. Stephan, K. Tiwari, J. Weiser, A. Wright, M. Gillespie, G. Wu, L. Stein, H. Hermjakob, P. D’Eustachio, The Reactome Knowledgebase 2026. Nucleic Acids Res. 54, D673–D681 (2026).

32. L. B. Norbom, B. Syed, R. Kjelkenes, J. Rokicki, A. Beauchamp, S. Nerland, A. Kushki, E. Anagnostou, P. Arnold, J. Crosbie, E. Kelley, R. Nicolson, R. Schachar, M. J. Taylor, L. T. Westlye, C. K. Tamnes, J. P. Lerch, Probing Autism and ADHD subtypes using cortical signatures of the T1w/T2w-ratio and morphometry. NeuroImage Clin. 45, 103736 (2025).

33. D. Pinto, E. Delaby, D. Merico, M. Barbosa, A. Merikangas, L. Klei, B. Thiruvahindrapuram, X. Xu, R. Ziman, Z. Wang, J. A. S. Vorstman, A. Thompson, R. Regan, M. Pilorge, G. Pellecchia, A. T. Pagnamenta, B. Oliveira, C. R. Marshall, T. R. Magalhaes, J. K. Lowe, J. L. Howe, A. J. Griswold, J. Gilbert, E. Duketis, B. A. Dombroski, M. V. De Jonge, M. Cuccaro, E. L. Crawford, C. T. Correia, J. Conroy, I. C. Conceição, A. G. Chiocchetti, J. P. Casey, G. Cai, C. Cabrol, N. Bolshakova, E. Bacchelli, R. Anney, S. Gallinger, M. Cotterchio, G. Casey, L. Zwaigenbaum, K. Wittemeyer, K. Wing, S. Wallace, H. van Engeland, A. Tryfon, S. Thomson, L. Soorya, B. Rogé, W. Roberts, F. Poustka, S. Mouga, N. Minshew, L. A. McInnes, S. G. McGrew, C. Lord, M. Leboyer, A. S. Le Couteur, A. Kolevzon, P. Jiménez González, S. Jacob, R. Holt, S. Guter, J. Green, A. Green, C. Gillberg, B. A. Fernandez, F. Duque, R. Delorme, G. Dawson, P. Chaste, C. Café, S. Brennan, T. Bourgeron, P. F. Bolton, S. Bölte, R. Bernier, G. Baird, A. J. Bailey, E. Anagnostou, J. Almeida, E. M. Wijsman, V. J. Vieland, A. M. Vicente, G. D. Schellenberg, M. Pericak-Vance, A. D. Paterson, J. R. Parr, G. Oliveira, J. I. Nurnberger, A. P. Monaco, E. Maestrini, S. M. Klauck, H. Hakonarson, J. L. Haines, D. H. Geschwind, C. M. Freitag, S. E. Folstein, S. Ennis, H. Coon, A. Battaglia, P. Szatmari, J. S. Sutcliffe, J. Hallmayer, M. Gill, E. H. Cook, J. D. Buxbaum, B. Devlin, L. Gallagher, C. Betancur, S. W. Scherer, Convergence of genes and cellular pathways dysregulated in autism spectrum disorders. Am. J. Hum. Genet. 94, 677–694 (2014).

34. S. De Rubeis, X. He, A. P. Goldberg, C. S. Poultney, K. Samocha, A. E. Cicek, Y. Kou, L. Liu, M. Fromer, S. Walker, T. Singh, L. Klei, J. Kosmicki, F. Shih-Chen, B. Aleksic, M. Biscaldi, P. F. Bolton, J. M. Brownfeld, J. Cai, N. G. Campbell, A. Carracedo, M. H. Chahrour, A. G. Chiocchetti, H. Coon, E. L. Crawford, S. R. Curran, G. Dawson, E. Duketis, B. A. Fernandez, L. Gallagher, E. Geller, S. J. Guter, R. S. Hill, J. Ionita-Laza, P. Jimenz Gonzalez, H. Kilpinen, S. M. Klauck, A. Kolevzon, I. Lee, I. Lei, J. Lei, T. Lehtimäki, C.-F. Lin, A. Ma’ayan, C. R. Marshall, A. L. McInnes, B. Neale, M. J. Owen, N. Ozaki, M. Parellada, J. R. Parr, S. Purcell, K. Puura, D. Rajagopalan, K. Rehnström, A. Reichenberg, A. Sabo, M. Sachse, S. J. Sanders, C. Schafer, M. Schulte-Rüther, D. Skuse, C. Stevens, P. Szatmari, K. Tammimies, O. Valladares, A. Voran, W. Li-San, L. A. Weiss, A. J. Willsey, T. W. Yu, R. K. C. Yuen, DDD Study, Homozygosity Mapping Collaborative for Autism, UK10K Consortium, E. H. Cook, C. M. Freitag, M. Gill, C. M. Hultman, T. Lehner, A. Palotie, G. D. Schellenberg, P. Sklar, M. W. State, J. S. Sutcliffe, C. A. Walsh, S. W. Scherer, M. E. Zwick, J. C. Barett, D. J. Cutler, K. Roeder, B. Devlin, M. J. Daly, J. D. Buxbaum, Synaptic, transcriptional and chromatin genes disrupted in autism. Nature 515, 209–215 (2014).

35. L. de la Torre-Ubieta, H. Won, J. L. Stein, D. H. Geschwind, Advancing the understanding of autism disease mechanisms through genetics. Nat. Med. 22, 345–361 (2016).

36. S. J. Sanders, M. T. Murtha, A. R. Gupta, J. D. Murdoch, M. J. Raubeson, A. J. Willsey, A. G. Ercan-Sencicek, N. M. DiLullo, N. N. Parikshak, J. L. Stein, M. F. Walker, G. T. Ober, N. A. Teran, Y. Song, P. El-Fishawy, R. C. Murtha, M. Choi, J. D. Overton, R. D. Bjornson, N. J. Carriero, K. A. Meyer, K. Bilguvar, S. M. Mane, N. Sestan, R. P. Lifton, M. Günel, K. Roeder, D. H. Geschwind, B. Devlin, M. W. State, De novo mutations revealed by whole-exome sequencing are strongly associated with autism. Nature 485, 237–241 (2012).

37. S. Marek, B. Tervo-Clemmens, F. J. Calabro, D. F. Montez, B. P. Kay, A. S. Hatoum, M. R. Donohue, W. Foran, R. L. Miller, T. J. Hendrickson, S. M. Malone, S. Kandala, E. Feczko, O. Miranda-Dominguez, A. M. Graham, E. A. Earl, A. J. Perrone, M. Cordova, O. Doyle, L. A. Moore, G. M. Conan, J. Uriarte, K. Snider, B. J. Lynch, J. C. Wilgenbusch, T. Pengo, A. Tam, J. Chen, D. J. Newbold, A. Zheng, N. A. Seider, A. N. Van, A. Metoki, R. J. Chauvin, T. O. Laumann, D. J. Greene, S. E. Petersen, H. Garavan, W. K. Thompson, T. E. Nichols, B. T. T. Yeo, D. M. Barch, B. Luna, D. A. Fair, N. U. F. Dosenbach, Reproducible brain-wide association studies require thousands of individuals. Nature 603, 654–660 (2022).

38. R. B. Mars, S. Jbabdi, M. F. S. Rushworth, A Common Space Approach to Comparative Neuroscience. Annu. Rev. Neurosci. 44, 69–86 (2021).

39. J. P. Lerch, A. J. W. Van Der Kouwe, A. Raznahan, T. Paus, H. Johansen-Berg, K. L. Miller, S. M. Smith, B. Fischl, S. N. Sotiropoulos, Studying neuroanatomy using MRI. Nat. Neurosci. 20, 314–326 (2017).

40. L. A. Schwarz, C. P. Dotter, S. Isaev, M. Lisi, D. Malzl, C. Büschl, S. Ladstätter, B. Oliveira, M. Barel, B. Basilico, C. Chintaluri, S. Gorkiewicz, M. Goudarzi, T. Belinova, S. Reichl, G. Sendžikaitė, S. A. Jayaram, P. Koppensteiner, C. Sommer, T. P. Vogels, J. Menche, I. Adameyko, P. V. Kharchenko, C. Bock, G. Novarino, Cortical development dynamics across autism spectrum disorder mouse models. Nature, doi: 10.1038/s41586-026-10679-1 (2026).

41. J. P. Lerch, J. G. Sled, R. M. Henkelman, MRI phenotyping of genetically altered mice. Magn. Reson. Neuroimaging Methods Protoc., 349–361 (2011).

42. L. S. Cahill, C. L. Laliberté, J. Ellegood, S. Spring, J. A. Gleave, M. C. van Eede, J. P. Lerch, R. M. Henkelman, Preparation of fixed mouse brains for MRI. Neuroimage 60, 933–939 (2012).

43. B. J. Nieman, N. A. Bock, J. Bishop, X. J. Chen, J. G. Sled, J. Rossant, R. M. Henkelman, Magnetic resonance imaging for detection and analysis of mouse phenotypes. NMR Biomed. Int. J. Devoted Dev. Appl. Magn. Reson. Vivo 18, 447–468 (2005).

44. B. J. Nieman, N. A. Bock, J. Bishop, J. G. Sled, X. Josette Chen, R. Mark Henkelman, Fast spin-echo for multiple mouse magnetic resonance phenotyping. Magn. Reson. Med. Off. J. Int. Soc. Magn. Reson. Med. 54, 532–537 (2005).

45. J. Dazai, S. Spring, L. S. Cahill, R. M. Henkelman, Multiple-mouse neuroanatomical magnetic resonance imaging. JoVE J. Vis. Exp., e2497 (2011).

46. J. Dazai, N. A. Bock, B. J. Nieman, L. M. Davidson, R. M. Henkelman, X. J. Chen, Multiple mouse biological loading and monitoring system for MRI. Magn. Reson. Med. Off. J. Int. Soc. Magn. Reson. Med. 52, 709–715 (2004).

47. T. L. Spencer Noakes, R. M. Henkelman, B. J. Nieman, Partitioning k-space for cylindrical three-dimensional rapid acquisition with relaxation enhancement imaging in the mouse brain. NMR Biomed. 30 (2017).

48. D. L. Collins, P. Neelin, T. M. Peters, A. C. Evans, Automatic 3D intersubject registration of MR volumetric data in standardized Talairach space. J. Comput. Assist. Tomogr. 18, 192–205 (1994).

49. B. B. Avants, C. L. Epstein, M. Grossman, J. C. Gee, Symmetric diffeomorphic image registration with cross-correlation: evaluating automated labeling of elderly and neurodegenerative brain. Med. Image Anal. 12, 26–41 (2008).

50. B. B. Avants, N. J. Tustison, G. Song, P. A. Cook, A. Klein, J. C. Gee, A reproducible evaluation of ANTs similarity metric performance in brain image registration. Neuroimage 54, 2033–2044 (2011).

51. M. Friedel, M. C. van Eede, J. Pipitone, M. M. Chakravarty, J. P. Lerch, Pydpiper: a flexible toolkit for constructing novel registration pipelines. *Front*. Neuroinformatics 8, 67 (2014).

52. A. N. Zizzo, L. Erdman, B. M. Feldman, A. Goldenberg, Similarity network fusion: a novel application to making clinical diagnoses. Rheum. Dis. Clin. 44, 285–293 (2018).

53. B. Wang, A. M. Mezlini, F. Demir, M. Fiume, Z. Tu, M. Brudno, B. Haibe-Kains, A. Goldenberg, Similarity network fusion for aggregating data types on a genomic scale. Nat. Methods 11, 333–337 (2014).

54. C. Von Mering, L. J. Jensen, B. Snel, S. D. Hooper, M. Krupp, M. Foglierini, N. Jouffre, M. A. Huynen, P. Bork, STRING: known and predicted protein–protein associations, integrated and transferred across organisms. Nucleic Acids Res. 33, D433–D437 (2005).

55. B. Jassal, L. Matthews, G. Viteri, C. Gong, P. Lorente, A. Fabregat, K. Sidiropoulos, J. Cook, M. Gillespie, R. Haw, The reactome pathway knowledgebase. Nucleic Acids Res. 48, D498–D503 (2020).

56. Y. Benjamini, Y. Hochberg, Controlling the false discovery rate: a practical and powerful approach to multiple testing. J. R. Stat. Soc. Ser. B Methodol. 57, 289–300 (1995).

57. V. Hus, C. Lord, The autism diagnostic observation schedule, module 4: revised algorithm and standardized severity scores. J. Autism Dev. Disord. 44, 1996–2012 (2014).

58. A. Ickowicz, R. J. Schachar, R. Sugarman, S. X. Chen, C. Millette, L. Cook, The parent interview for child symptoms: a situation-specific clinical research interview for attention-deficit hyperactivity and related disorders. Can. J. Psychiatry 51, 325–328 (2006).

59. S. A. Bedford, M. T. M. Park, G. A. Devenyi, S. Tullo, J. Germann, R. Patel, E. Anagnostou, S. Baron-Cohen, E. T. Bullmore, L. R. Chura, Large-scale analyses of the relationship between sex, age and intelligence quotient heterogeneity and cortical morphometry in autism spectrum disorder. Mol. Psychiatry 25, 614–628 (2020).

60. M. J. Hawrylycz, E. S. Lein, A. L. Guillozet-Bongaarts, E. H. Shen, L. Ng, J. A. Miller, L. N. Van De Lagemaat, K. A. Smith, A. Ebbert, Z. L. Riley, An anatomically comprehensive atlas of the adult human brain transcriptome. Nature 489, 391–399 (2012).

61. E. S. Lein, M. J. Hawrylycz, N. Ao, M. Ayres, A. Bensinger, A. Bernard, A. F. Boe, M. S. Boguski, K. S. Brockway, E. J. Byrnes, Genome-wide atlas of gene expression in the adult mouse brain. Nature 445, 168–176 (2007).

62. Q. Wang, S.-L. Ding, Y. Li, J. Royall, D. Feng, P. Lesnar, N. Graddis, M. Naeemi, B. Facer, A. Ho, The Allen mouse brain common coordinate framework: a 3D reference atlas. Cell 181, 936–953. e20 (2020).

